# Ascertaining cells’ synaptic connections and RNA expression simultaneously with massively barcoded rabies virus libraries

**DOI:** 10.1101/2021.09.06.459177

**Authors:** Arpiar Saunders, Kee Wui Huang, Cassandra Vondrak, Christina Hughes, Karina Smolyar, Harsha Sen, Adrienne C. Philson, James Nemesh, Alec Wysoker, Seva Kashin, Bernardo L. Sabatini, Steven A. McCarroll

## Abstract

Brain function depends on forming and maintaining connections between neurons of specific types, ensuring neural function while allowing the plasticity necessary for cellular and behavioral dynamics. However, systematic descriptions of how brain cell types organize into synaptic networks and which molecules instruct these relationships are not readily available. Here, we introduce SBARRO (<u>S</u>ynaptic <u>B</u>arcode <u>A</u>nalysis by <u>R</u>etrograde <u>R</u>abies Read<u>O</u>ut), a method that uses single-cell RNA sequencing to reveal directional, monosynaptic relationships based on the paths of a barcoded rabies virus from its “starter” postsynaptic cell to that cell’s presynaptic partners^1^. Thousands of these partner relationships can be ascertained in a single experiment, alongside genome-wide RNA profiles – and thus cell identities and molecular states – of each host cell. We used SBARRO to describe synaptic networks formed by diverse mouse brain cell types *in vitro*, leveraging a system similar to those used to identify synaptogenic molecules. We found that the molecular identity (cell type/subtype) of the starter cell predicted the number and types of cells that had synapsed onto it. Rabies transmission tended to occur into cells with RNA-expression signatures related to developmental maturation and synaptic transmission. The estimated size of a cell’s presynaptic network, relative to that of other cells of the same type, associated with increased expression of *Arpp21* and *Cdh13*. By tracking individual virions and their clonal progeny as they travel among host cells, single-cell, single-virion genomic technologies offer new opportunities to map the synaptic organization of neural circuits in health and disease.

## MAIN

The mammalian brain contains hundreds of cell types that connect with one another through synapses into intricate, and mostly uncharacterized, neural circuits. Traditional approaches for measuring synaptic connections and networks – such as whole-cell electrophysiological recordings and anatomical reconstructions from electron microscopy – sample only a few cells or small tissue volumes, do not readily scale to many animals or genotypes, and do not ascertain the molecular type and state of each cell. Recent advances in single-cell transcriptomic profiling have made identifying cells and cell types within complex tissue routine^2–5^. Together with engineered proteins and viruses, additional cell features such as protein expression^2^, developmental origin^3, 4^, axonal projection patterns^5^ and physical interactions^6^ can be decoded from RNA data. In the nervous system, rabies virus spreads from cell to cell in a retrograde fashion, from a neuron’s dendrites into the axons of its presynaptic partners^7^. Prevailing models suggest such transmission events occur at synapses, likely due to the presence of viral entry receptors^8^ and high rates of membrane turnover. While the synaptic phenomenology of rabies virus transmission has been used for decades to discover neural pathways^9^, inefficient conversion of plasmid DNA into infective RNA-containing particles has largely precluded using rabies and other *lyssaviruses* in genomic applications^10^.

Here, we introduce SBARRO (<u>S</u>ynaptic <u>B</u>arcode <u>A</u>nalysis by <u>R</u>etrograde <u>R</u>abies Read<u>O</u>ut), which combines monosynaptic rabies virus tracing, viral genomic barcoding and scRNA-seq to generate high-throughput descriptions of cell-type-resolved synaptic networks. In SBARRO, encapsidated rabies virus genomes are distinguished by unique, transcribed viral barcode sequences (VBCs), allowing thousands of monosynaptic networks to be reconstructed in parallel by tracking paths of clonal infections which originate in postsynaptic starter cells and spread to those cells’ presynaptic partners. By replacing the endogenous glycoprotein gene (*G*) – necessary for viral spread – with *EGFP* in the viral genome, rabies virus transmission is restricted to cells that are directly presynaptic^1, 11^. In sampling cellular RNAs alongside VBCs, our approach reveals: 1) postsynaptic vs presynaptic cell identities; 2) cell types and molecular states (via host cell RNAs); and synaptic networks (via shared VBCs; Fig. 1a). In genomics, rabies virus has been previously used in RABID-seq^6^, in which rabies virus spread was used to infer putative direct physical contacts between glia, a previously unknown and uncharacterized type of rabies virus transmission.

**Figure 1.**
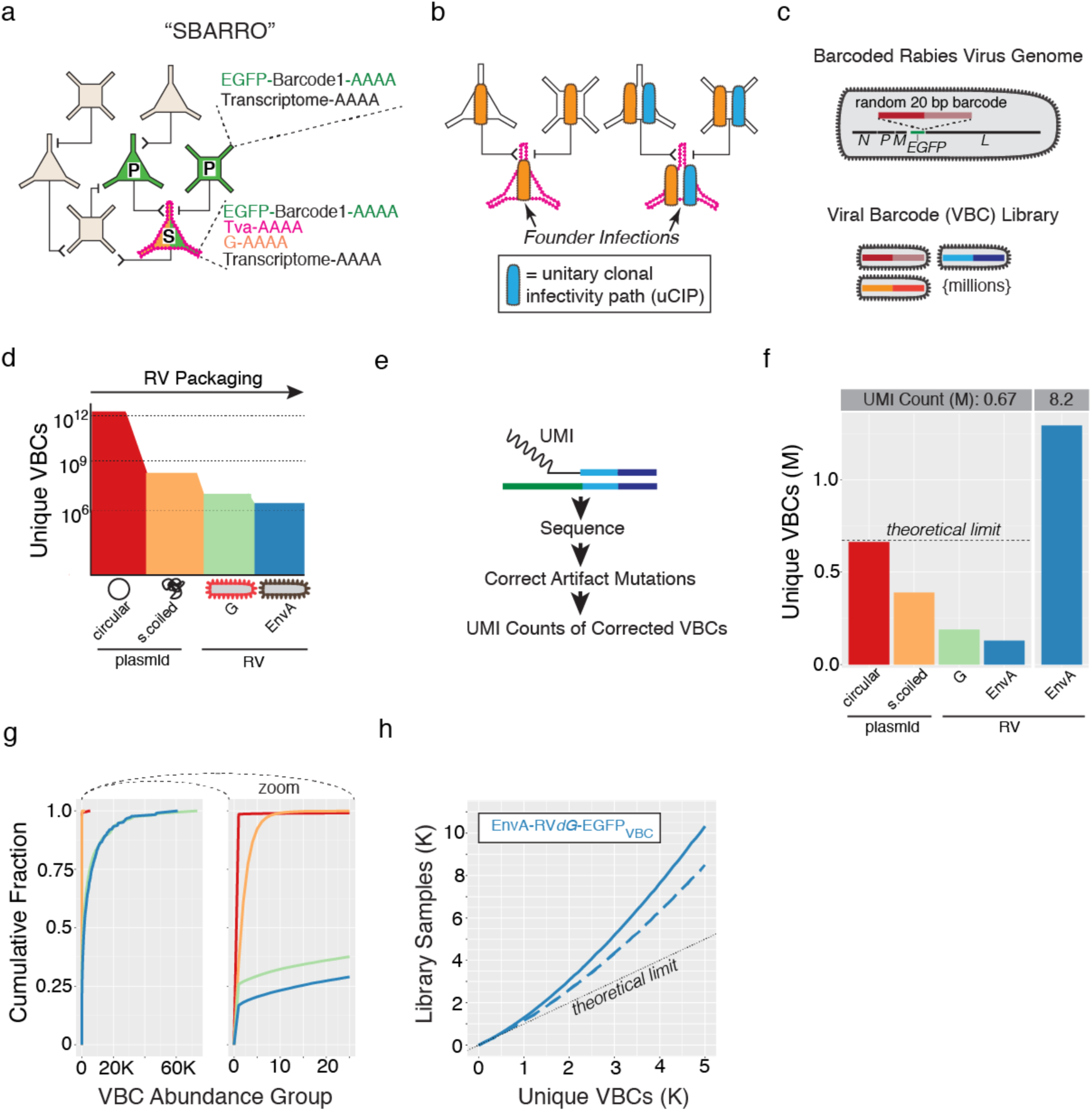
Single-virion RNA tracking enabled by libraries of rabies virus particles encapsidating millions of uniquely barcoded genomes. **a.** Monosynaptic SBARRO schematic. TVA-expressing starter cells (“S”) complemented in *trans* with rabies virus glycoprotein (G) are selectively transduced by EnvA-pseudotyped rabies virus in which *G* has been replaced barcoded *EGFP* (EnvA-RV*dG*-*EGFP*VBC). G-complemented clonal particles spread a single retrograde synapse into presynaptic partner cells (“P”). Single-cell RNA profiles inform 1) synaptic groupings (from *EGFP* based viral barcode (VBC) sharing); 2) Starter or presynaptic status (from *TVA* mRNAs) and 3) host cell type (by capturing thousands of cellular mRNAs). **b.** Monosynaptic relationships are inferred through unitary clonal infectivity paths (uCIPs), defined by a subset of VBCs carried by those rabies virus particles sufficiently rare enough in the infecting library to seed single founder infections in starter cells. **c.** Rabies virus particles are distinguished from each other by a 20 bp bipartite VBC in the 3’ UTR of *EGFP*. VBC libraries contain millions of unique particles. **d.** Schematic of VBC diversity during barcoded rabies virus packaging. **e.** Schematic of sequencing-based genomic VBC quantification using unique molecular identifiers (UMIs; Extended Data Fig. 1c). Post-cellular polymerase mutations incurred during library amplification and sequencing were corrected informatically (Extended Data Fig. 2b and **Methods**). **f-h.** VBC diversity metrics (color-coded as in panel **d**). **f.** Unique VBCs identified by 0.67 million (M) UMI counts across each packaging stage (*left*) or with ∼12.2 fold more counts (8.2 M) after EnvA pseudotyping (*right*). **g.** Cumulative distribution of VBCs binned by “VBC abundance group” (AG) across packaging stages (0.67 M counts / stage). Total counts for all VBCs sampled once belong to AG = 1; sampled twice belong AG = 2, etc. **h.** The relationship between the number of unique VBCs identified after a given number of *in silico* samples drawn from the EnvA-RV*dG*-*EGFP*VBC library (8.2 M counts; blue line) and after removing the 88 most abundant VBCs (dashed blue line). The dotted line shows maximum theoretical diversity (in which every drawn VBC is unique).

Here we present a comprehensive experimental and analytical framework for using SBARRO to discover structural and molecular properties of cell-type-specific synaptic connectivity. We leverage dissociated mouse brain cells grown into synaptic networks *in vitro*, similar to systems used to identify synaptogenic molecules^12, 13^ and for which features of *in vivo* connectivity can remain^14^. We discover that presynaptic network properties, such as cell type composition and size, are conferred in part by the postsynaptic cell type. Finally, we discover that rabies virus spread associates with a molecular signature of synaptic maturation, suggesting that functioning synapses, in addition to entry receptors, are critical for uptake of rabies virus.

### Barcoding millions of rabies viral genomes

Synaptic tracing with barcoded rabies requires libraries of barcoded rabies virus particles that have high numbers of unique barcodes which are as uniform as possible in abundance, such that individual viral barcodes are introduced into no more than one starter cell (in an experiment) and can be used to define “unitary” clonal infectivity paths (uCIPs; Fig. 1b). Inefficiencies in creating negative-stranded RNA viruses from DNA have historically precluded generating complex rabies libraries^10^. To generate libraries encoding millions of barcodes, we developed molecular and computational methods to introduce, retain, and quantify barcodes in DNA plasmids and in rescued RNA genomes (Fig. 1c,d; Extended Data Fig. 1 and Extended Data Fig. 2).

We first developed a PCR-based strategy to flexibly engineer bipartite barcodes, generated through combinatorial diversity, into circular DNA plasmids (Extended Data Fig. 1b), followed by transformation and plate-based growth conditions optimized to retain DNA plasmid barcode diversity (Extended Data Fig. 2b,c). We also created a rabies rescue system – achieving equivalent viral titers 3-fold faster than the current protocols^15^ – that minimized barcode loss and disproportionate amplification during viral replication (Extended Data Fig. 2d). To assess viral barcode diversity and distribution, we used single-molecule sequencing (Fig. 1e and Extended Data Fig. 1c), for which we developed analysis methods to identify and correct for PCR and sequencing mutations (Extended Data Fig. 2b and **Methods**).

We used this approach to generate an EnvA-pseudotyped rabies library with a 20 bp randomer encoded in the 3’ UTR of *EGFP* of SAD-dG-B19 (EnvA-RV*dG*-*EGFP*VBC; Extended Data Fig. 1a). We compared the total number of unique barcodes and their relative abundances across each production stage (Fig. 1d-g and Extended Data Fig. 1a). After PCR and circularization, nearly every sequenced plasmid contained a unique barcode. This diversity was reduced by bacterial amplification, though without substantially distorting representation of the retained barcodes. Rabies rescue induced barcode loss and abundance distortions and was mildly exacerbated by EnvA- pseudotyping. Deeper sequencing of the final EnvA-RV*dG*-*EGFP*VBC genomes (6.4 unique molecular identifiers (UMI) per viral barcode on average) quantified the relative abundances of 1.29 million unique, error-corrected barcodes.

To estimate the fraction of EnvA-RV*dG*-*EGFP*VBC founder infections that would be from viral particles with unique barcodes, we performed *in silico* mock infections by randomly sampling barcodes from the sequenced genomes of the infecting library and calculated the resulting number of unique barcodes (Fig. 1h). For these analyses, we used 50% unique barcodes as our benchmark, though the actual number of unique founder infections depends on properties of the infecting library and the number of founder infections in the experiment. Sampling up to ∼8,900 library genomes resulted in >50% unique barcodes; this could be increased to 15,500 library genomes by filtering out the 88 most abundant barcodes in the library, and to 83,600 library genomes by mixing 9 equivalent libraries *in silico* (Extended Data Fig. 2e). These analyses suggested that our optimized protocols have helped overcome inefficiencies that previously limited rabies applications in scalable genomics research and suggested uCIPs can be efficiently generated from thousands of founder infections.

### Characterizing the barcoded rabies library with 28,000 founder infections

To directly determine the relationship between our barcoded EnvA-RV*dG*-*EGFP*VBC particles and the brain cells they infect, we infected three replicate cell cultures derived from embryonic mouse cortex. Infections were targeted to cells by recombinant adeno-associated virus (rAAV) expression of TVA; the host cells lacked the G protein necessary for rabies spread (Fig. 2a,b and **Methods)**. After 72 hours, we collected 60,816 transcriptomes (n=6 scRNA-seq libraries each from a single culture well) that captured both the cellular RNAs and the barcoded region of *EGFP* mRNAs (Extended Data Fig. 3 and **Methods)**.

**Figure 2.**
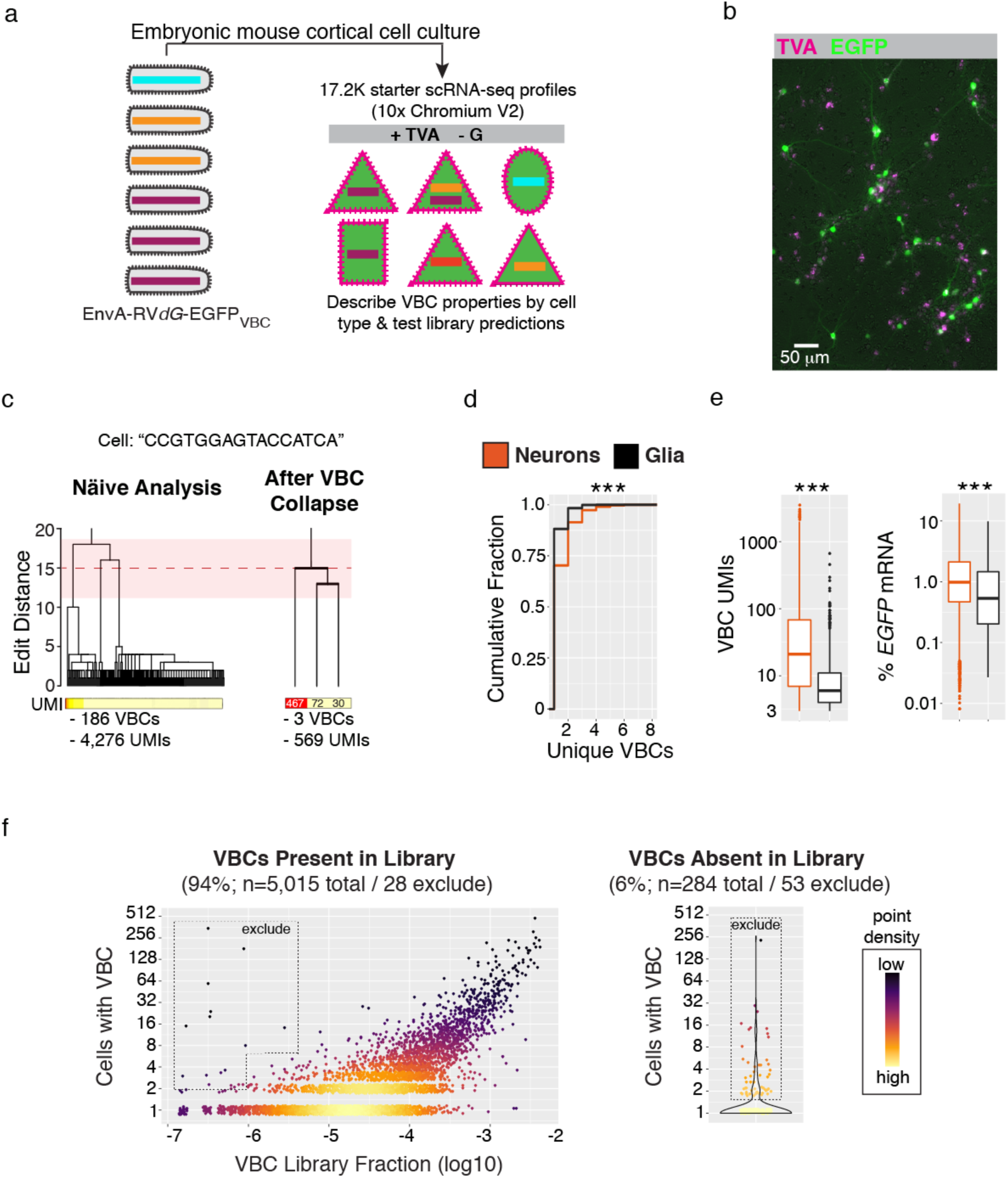
Properties of barcoded library infection revealed through single-cell RNA profiling from mouse brain cultures lacking cell-to-cell viral spread. **a.** Experimental schematic. The EnvA-RV*dG*-*EGFP*VBC library transduced starter cells (+TVA) from which the rabies virus could replicate but not spread to other cells (-G). RNA profiles were captured (n=60.8K cells), including from cells infected by rabies virus which function as a corpus of starter cells (n=17.2K). **b.** Representative image of dissociated mouse brain cell cultures (14 days *in vitro*) expressing TVA (magenta) and EGFP (green). Cultures consist of TVA-/EGFP-, TVA+/EGFP- and TVA+/EGFP+ cells. **c.** Inference of founder VBC sequences and accurate UMI-based counts from single cell RNA profiles in light of subsequent barcode mutations. A dendrograms illustrating VBC sequence relationships *(top*) and UMI counts (*below*) before (*left*) and after (*right*) “within-cell VBC collapse” for a single example RNA profile (**Methods**). The mean (red dotted line) and two standard deviations (pink shading) from distribution of edit distances among random barcode sequences. **d-e**. Comparison of single cell VBC properties ascertained from RNA profiles of neurons (n= 4,222) or glia (n= 914; *** = p < 2.2e-16, Kolmogorov–Smirnov Test). Only data from 1:10 EnvA-RV*dG*-*EGFP*VBC dilution are shown (**Methods** and Extended Data Fig. 3f). **d**. Cumulative distribution of unique VBCs. **e**. Total VBC UMIs (*left*) or % EGFP mRNA (*right*). **f.** Critically evaluating the performance of the EnvA- RV*dG*-*EGFP*VBC through a corpus of 17.2K starter cell RNA profiles. *Left*, for ascertained VBCs in the library (94%), the relationship between library abundance and the number of independent starter cell infections. *Right*, for library-absent VBCs (6%), the number of independent infections. VBCs observed in more starter cell RNA profiles than expected based on quantitative library abundance or library absence were flagged for exclusion (**Methods**).

Naïve analysis of barcode sequences from individual cells initially suggested large barcode “families” with many highly similar sequences (Fig. 2c). Reasoning that such relationships were largely created by PCR or sequencing errors, we developed an algorithm to collapse families of highly similar barcode sequences into the single barcode responsible for the putative founder infection (**Methods**). After collapse, barcode sequences associated with different inferred founder infections in the same cells had the same distribution of similarity relationships (edit distances) as random barcodes did. Furthermore, there were (1) similar numbers of UMIs covering the barcode and *EGFP* transcript within the same cells and (2) independence in the number of unique barcodes and barcode-associated RNA counts (Extended Data Fig. 3a,b). All data presented hereafter have been computationally collapsed in this way.

Cells in which we detected at least one viral barcode also tended to have devoted a substantial fraction of their transcription to rabies genes (% of total UMIs mean±sem, 15±0.14), compared to cells that did not have a barcode (0.3±0.007%, a rate consistent with background due to ambient cell-free RNA). This suggests that barcode ascertainment was sensitive, selective, and distinguished infected starter cells (n=17,283) from neighboring uninfected (n= 43,533) cells (Extended Data Fig. 3c). Putatively infected cells (those cells for which >1% of total UMIs came from rabies genes) for which we failed to detect a barcode tended to have very small RNA profiles (< 500 UMIs). Additionally, less than 2% of cells with a barcode and a large RNA profile (>10,000 UMIs) lacked viral loads indicative of infection (<1% rabies RNA), suggesting spurious viral barcode associations were rare.

Using the above data, we investigated whether the properties of infection differed among starter cell types. We found that infection mainly occurred in glutamatergic neurons, interneurons and astrocytes, and was less frequently observed in other glia types (polydendrocytes and oligodendrocytes), neural precursor cells, and cells undergoing mitosis (Extended Data Fig. 3e and **Methods)**. Even among infected cells, analysis revealed clear differences in infection properties: relative to infected glia, infected neurons tended to have more founder infections (unique barcodes mean±sem: Neurons, 1.4±0.01; Glia, 1.1±0.01), far more barcoded rabies transcripts detected per RNA profile (Neurons, 97.7±4.0; Glia, 13.8± 1.1), and higher percentages of *EGFP* per RNA profile (Neurons, 1.5±0.03; Glia, 1.1±0.05)(Fig. 2d,e), revealing previously unknown cell-type-specific properties of rabies virus infection.

In principle, the combination of multiple founder infections in the same starter cell could help define uCIPs through coupled presynaptic spread, but in practice, cell biological constraints might limit the number of founder infections. To evaluate this, we leveraged the viral barcodes to quantify the multiplicity of infection (MOI) at single-cell resolution and to relate this to the titer of the infecting library (Extended Data Fig. 3f). At the lowest titer we tried (MOI, ∼0.15), more than 97% of neuron and astrocyte RNA profiles were associated with a single VBC (unique VBCs mean±sem for neurons/astrocytes: MOI ∼0.15, 1.07±0.009/1.03±0.01). Infections with 10-fold higher titer resulted in more multiply infected cells (with two or more viral barcodes)(MOI ∼1.5, 1.42±0.01/1.14±0.01). However, we saw only minimal further increases at 100-fold higher titer (MOI ∼15, 1.6±0.01 / 1.2±0.02). At a biological level, these data suggest an intrinsic asymptote in the number of founder infections individual cells will meaningfully sustain – perhaps, for example, because cell-biological machinery are effectively hijacked by the earliest founders. At an engineering level, these results also suggested rabies virus titer-ranges for efficiently transducing starter cells with multiple viral barcodes.

Infecting and analyzing large numbers of starter cells in these control, no-spread experiments helped us to better understand many properties of rabies virus infections and barcoded rabies libraries. We compared the EnvA-RV*dG*-*EGFP*VBC library abundances of 1.29 million barcodes to their 28,755 founder infections distributed across 17,283 starter cells (Fig. 2f). Critically for later inferences, the abundance of a barcode in the infecting library predicted the number of cells it would infect (Fig. 2f). (A few barcodes that appeared to overperform this expectation were flagged for computational removal from future analyses, Fig. 2f). In addition, some 6% of infections involved viral barcodes that we had not detected by sequencing the library, presumably because they were present at very low abundance (Fig. 2f) (any of these that infected multiple cells in this “no-spread” experiment were also flagged for removal from future analyses). Intriguingly, a small number of barcode pairs consistently appeared together in the same starter cells, even in distinct experiments. Because rabies particles do not have strict genome size limitations^10, 16^, we reasoned that barcode interdependence might result from concatenated genomes. (These pairs were also flagged and removed from future analyses; **Methods**). These analyses suggest the abundance of barcodes in library genomes has considerable predictive power in estimating the number of starter cell founder infections, but also highlight examples in which individual barcodes or barcode pairs defy expectations. Thus, each SBARRO infecting library should be carefully evaluated in a large number of starter cells, as we describe further below.

### Massively parallel inference of monosynaptic relationships between cells

We next sought to describe cell-type-specific synaptic wiring of an *in vitro* culture. We focused on *in vitro* experiments because such systems 1) have been used extensively to screen for genes and molecules involved in synapse development; 2) can retain features of cell-type-specific connectivity, and 3) have more easily recoverable cells; in our hands, recovery of rabies infected neurons after *in vivo* experiments was inefficient. To increase cell type diversity, we co-cultured cells dissociated from embryonic cortex, striatum and caudal olfactory areas. We sparsely seeded potential starter cells in each culture well by using rAAVs to express TVA and the rabies glycoprotein (G), thus enabling EnvA-mediated rabies founder infections and G-dependent presynaptic spread. (Sparsity minimizes the opportunity for starter cells to become secondarily infected as presynaptic cells, which could in principle support polysynaptic spread). Lastly, after 12 days *in vitro*, during a period of prolific synaptogenesis^17^, starter cells were transduced with EnvA-RV*dG*-*EGFP*VBC (MOI, ∼1.5)(Fig. 3a and **Methods**). After another 72 hours, EGFP fluorescence was observed in putative presynaptic cells, many of which were spatially clustered around each dual-labeled starter cell and were interspersed with large numbers of uninfected cells (Extended Data Fig. 4a,b). This spatial pattern of rabies spread was consistent with the idea that the probability of neuronal connectivity scales roughly with spatial proximity and suggested that presynaptic networks innervating distinct starter cells were largely non-overlapping. No transduction (EGFP+ cells) was observed without TVA receptor expression, suggesting that all infections entered experiments through starter cells (data not shown).

**Figure 3.**
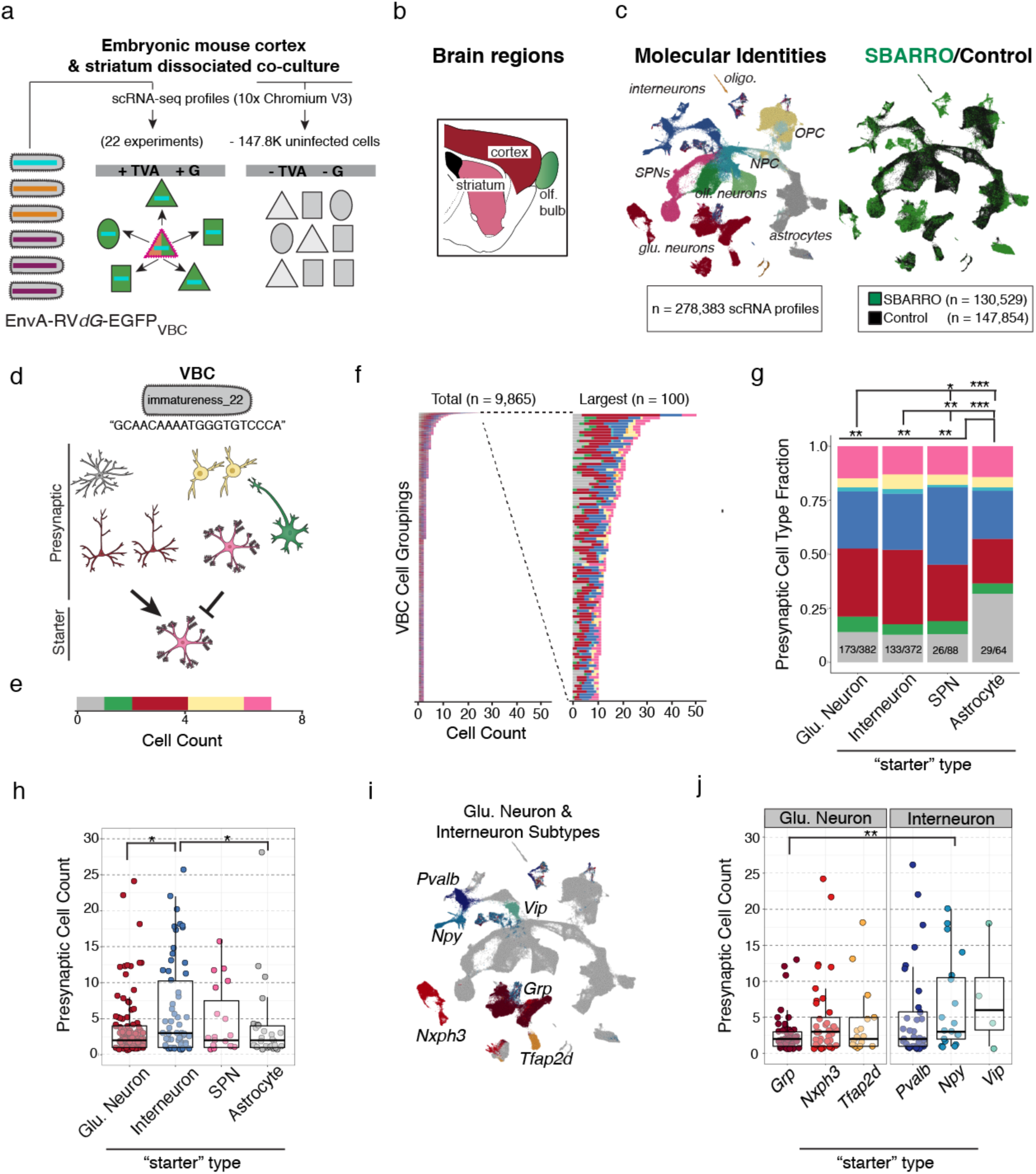
Massively parallel inference of cell-type-specific synaptic connectivity using SBARRO. **a.** Experimental schematic. The EnvA-RV*dG*-*EGFP*VBC library transduced starter cells (+TVA/+G) from which individual virion clonally replicate and undergo retrograde, monosynaptic spread into presynaptic cells. scRNA-seq libraries were prepared from either 1) SBARRO EGFP+ cells (n=23 culture wells from n=3 mouse preparations; n=130.5K scRNA profiles) or 2) preparation-matched control cells (n=147.6K scRNA profiles). **b.** Sagittal mouse brain schematic color-coded by region from which cells were co-cultured. **c.** UMAP embedding of scRNA profiles color-coded and labeled by coarse molecular identity (*left,* Extended Data Fig. 5a) or SBARRO/control status (*right*) following LIGER analysis (**Methods**). **d.** An example SBARRO network inferred through shared expression of VBC assigned the named “immatureness_22”. (Names were assigned to each VBC to better track VBC identities; **Methods**). Rabies particles encapsidating the “immatureness_22” genome are rare enough such that transduction of more than one starter cell founder infection is estimated to have a < 1% chance of occurring (**Methods**). The “immatureness_22” network consists of a 7 scRNA profiles with associated molecular identities, including an SPN starter cell and heterogenous collection of putative presynaptic cells (color-coded as in **c**). **e.** Cell-type composition of the inferred “immatureness_22” presynaptic network represented as a horizontal bar plot. **f.** Horizontal bar plots for n=9,865 inferred SBARRO networks with >= 2 cells (*left*) and the largest 100 networks (*right*). Of all networks, n=365 networks (3.7%) included starter cell assignments. **g-i.** Properties of inferred presynaptic networks stratified by starter cell type. **g.** Fractional cell-type compositions from inferred presynaptic networks exhibit quantitative differences across starter cell types (* = p <0.05; ** = p < 0.01; *** = p <0.001, Chi-Square Test). The number of aggregated networks and total presynaptic cells are shown (networks/total cells). **h.** Inferred presynaptic network sizes differ by starter cell type. (** = p < 0.05, Wilcoxon Test). **i.** UMAP embedding color-coded and labeled by glutamatergic neuron and interneuron subtype (Extended Data Fig. 5a,b). **j.** Inferred presynaptic network sizes by starter cell subtype (** = p < 0.01; Wilcoxon Test).

We identified cell types in thousands of reconstructed monosynaptic networks, profiling RNA from EGFP+ cells from 23 distinct culture wells (n=3 cell culture replicates). To learn the molecular identities of each cell and determine in what ways the infected population might be different from the total ensemble of cultured cells, we co-clustered scRNA profiles from SBARRO experiments (n=130,529 cells; mean UMIs, 17,622) and uninfected control cells (n=147,854 cells; mean UMIs, 18,117) based on shared host-cell RNA signatures (**Methods**). Cultured cell RNA profiles were from diverse and developmentally dynamic cell populations (Fig. 3b,c). We identified four populations of glutamatergic neurons; polydendrocytes; oligodendrocytes; and neural precursor cells (NPCs) developing into astrocytes and several mature GABAergic lineages (including three major interneuron populations, two olfactory-related neuron types, and spiny projection neurons (SPNs); Extended Data Fig. 5a,b). Neuronal identity assignments were confirmed by an integrated analysis with scRNA profiles from adult mouse neocortex^18^ (Extended Data Fig. 5c). Compared to the relative abundance of control cells, rabies-infected cells were enriched among mature interneurons (log2(rv/control) = 1.21), SPNs (0.94), glutamatergic neurons (0.83) and astrocytes (0.32), and depleted from developmentally immature cells (NPCs = -2.45; immature neurons = -2.44), mature GABAergic olfactory types (−1.67), oligodendrocytes (−0.94) and polydendrocytes (- 0.91)(Extended Data Fig. 5d).

We detected putative synaptic networks as clonal expansion of viral barcodes observed across cells (Extended Data Fig. 6a). Paired anatomical/SBARRO datasets suggests that roughly 10% of infected cells entered our single-cell analyses; the missing cells were likely lost or destroyed during physical dissociation and FACS-enrichment or remained unsampled after microfluidics-based RNA barcoding. Thus, synaptic networks are detectable yet contain only a small subset of the cells associated with each network (Extended Data Fig. 4c,d).

We identified starter cells by their TVA expression (Extended Data Fig. 7a-d and **Methods**). Presynaptic and starter cells were composed of similar cell types, but starter cells expressed a larger number of unique barcodes (mean±sem: starter, 4.3±1; presynaptic, 2.8±0.007, p < 2.2e-16, Kolmogorov–Smirnov Test;Extended Data Fig. 7e,f), which is expected as some barcodes may fail to transit and infect other cells in the analysis. Comparing FACS-based counts of fluorescently labelled starter or presynaptic cells suggested that starter cells failed to enter our analyses more frequently than presynaptic cells did (2.5 vs 25%; Extended Data Fig. 4c,d); this could reflect increased fragility and loss due to prolonged infection, or insufficient ascertainment of recombined TVA mRNAs. Thus, we expect many of our identified synaptic networks to be “orphaned” from their starter cell.

We developed a statistical framework to filter one or more co-expressed viral barcodes based on the 1) estimated number of founder infections and the 2) barcode abundance in the infecting library (**Methods**). We also excluded barcodes (n=551) or barcode pairs (n=689) that (in the control experiments) infected or co-infected more cells than expected based on their library abundance (Fig. 2f; Extended Data Fig. 6b; **Methods**). For example, in an experiment estimated to contain 2,484 total founder infections, we observed one example barcode in an SPN starter cell and seven diverse presynaptic cells (Fig. 3d). Based on the low abundance of the barcode in the infecting library (frequency = 3.5×10^-6^), we estimate that this barcode had a <1% chance of participating in more than one founder infection in this experiment. Thus this barcode passed our threshold (of <10%) and defined a uCIP (**Methods**).

We retained n=1,810 of 5,142 total viral barcodes, which alone or in combination, enabled n=9,865 non-redundant uCIP inferences of synaptic networks with >= 2 cells (n = 21,458 scRNA profiles; Fig. 3f). Inferred networks contained 2-52 cells (mean = 3.1, median = 2), consistent with ∼10% ascertainment of rabies-infected cells in culture (Extended Data Fig. 4c,d). Inferred networks contained predominantly neurons (79%) of diverse types, with smaller contributions from astrocytes (15%) and polydendrocytes (5%).

To determine whether the cell-type composition or the size of presynaptic networks varied across postsynaptic cell types, we focused on the 3.7% of networks with an identified starter cell (n=365 of 9,865 total networks). Presynaptic cell types differed according to postsynaptic cell type, as pair-wise comparisons suggested quantitative differences in presynaptic cell-type proportions between astrocytes versus neurons (p = 0.007 - 0.001, Chi-Square Test) and glutamatergic neurons and interneurons versus SPNs (p = 0.01 and 0.0002), but not between glutamatergic neurons and interneurons (p > 0.05, Chi-Square Test; Fig. 3g).

The sizes of inferred presynaptic networks exhibited variance that was partially explained by postsynaptic cell type (p = 0.048, Kruskal-Wallis Test). Pair-wise comparisons revealed that glutamatergic neurons and astrocytes tended to have smaller presynaptic networks than interneurons did (p = 0.018 – 0.022, Wilcoxon Test), while SPN presynaptic networks did not detectably differ from those of other cell types (n=109 glutamatergic neurons, mean±sem presynaptic cells= 3.6±0.4; n=29 astrocytes, 3.7±1.02; n=61 interneurons, 6.7±1.0; n=12 SPNs, 4.6±1.1; Fig. 3h). Differences in neuronal presynaptic network size appeared to be driven in part by cell subtypes (Fig. 3i,j and Extended Data Fig. 6c,d); while neuronal subtype categories did not, as a whole, rise to predictive significance in explaining variance in presynaptic network size (p = 0.08, Kruskal-Wallis test), paired comparisons revealed that *Grp*+ glutamatergic neurons and *Npy*+ interneurons tended to have small (n=50, mean±sem = 2.7±0.37 cells) and large (n=20, mean = 8±1.95 cells) presynaptic networks (p = 0.004, Wilcoxon Test), respectively. These results indicate that the number and molecular composition of putative presynaptic cells in an inferred network are qualitatively similar across postsynaptic cell types at an early, promiscuous stage of synaptogenesis *in vitro*, but highlight important exceptions in which postsynaptic cell type biases the number and classes of putative presynaptic partner cells. Differences in the number of presynaptic partner cells might relate to dendritic size differences *in vivo*. For example, compared to adult mouse neocortex, *Grp*+ glutamatergic neurons are most similar to L2/3 IT and L4/5 IT subtypes found in superficial cortical layers, which tend to have small dendritic arbors, while *Nxph3+* glutamatergic neurons are most similar to L6b, L5 NP and L6 CT subtypes found in deeper cortical layers, which tend to have larger dendritic arbors^18, 19^ (Extended Data Fig. 5c).

### Postsynaptic RNAs associated with presynaptic network properties

The formation and selective stabilization of synapses is shaped by competitive processes driven by molecular variation within^20^ and across cell populations^21^, yet many of the molecules remain unknown and incompletely understood. We sought to use the data from these experiments – in which synaptic connectivity inferences and molecular properties were measured in the same cells – to analyze how molecular variation associated with the properties of cell-type-specific networks.

We first asked whether presynaptic network size was explained by infection magnitude or innate immune response in starter cells, since these properties of infection could skew the results (Fig. 4a). We separated starter cells into two groups based on presynaptic network size, ranging from networks of 2–4 ascertained cells (“small”) or 7–52 ascertained cells (“large”) and four groups based on starter cell type (Fig. 4b). We compared both the viral load (the fraction of total cellular mRNAs from the rabies virus genome) and an aggregate innate immunity expression score (n=564 genes^22^) across these groups (n=144 scRNA profiles; Extended Data Fig. 8a). We found that, while both infection metrics varied by starter cell type (viral load, p = 0.05; innate immune expression score, p = 5.2 x 10^-12^, Two-way ANOVA Test), they did not associate with the presynaptic network size (viral load, p = 0.10; innate immune expression score, p = 0.16).

**Figure 4.**
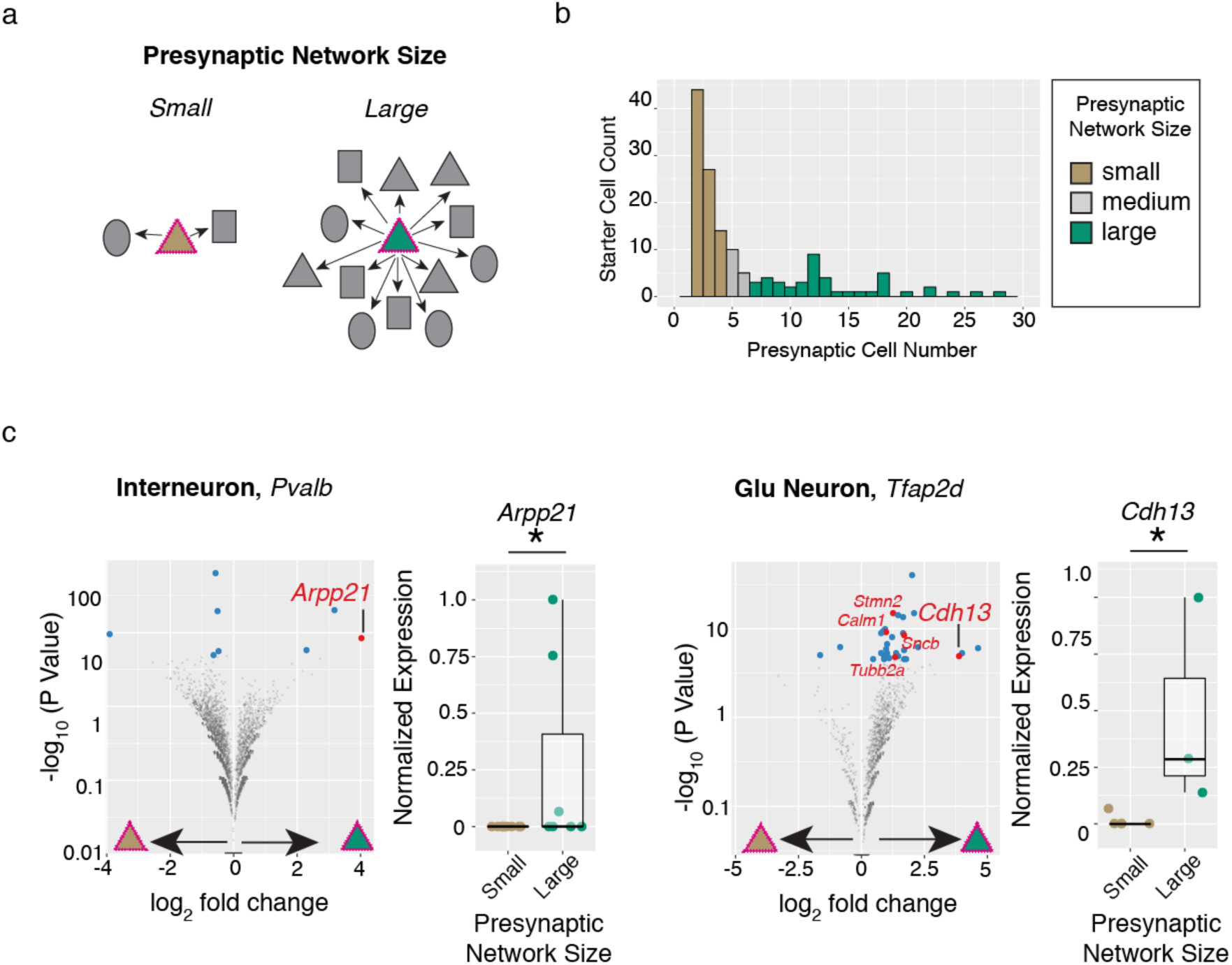
Postsynaptic RNA levels associated with rabies-based inferences of presynaptic network size. **a.** Schematic of postsynaptic starter cells with small (brown triangle) or large (green triangle) numbers of presynaptic partner cells. **b**. Histogram of inferred presynaptic network sizes for n=144 starter cell RNA profiles belonging to one of four major cell types (glutamatergic neurons, interneurons, SPNs or astrocytes). **c,d.** Differential expression testing identifies *Arpp21* upregulated in *Pvalb* interneurons (n=9 small versus n=7 large RNA profiles) and *Cdh13* as *Tfap2d* Glutamatergic neurons (n=5 small versus n=3 large RNA profiles) starter cells with large presynaptic networks. *Left*, Volcano plots illustrating results from differential expression testing of starter cell subtypes in which UMI counts were aggregated by inferred presynaptic network size category (Fisher’s Exact Test; **Methods**). Genes passing corrected p value thresholds (p < 0.05, blue dots) were further tested for differences in single-cell scaled expression (Wilcoxon Test; **Methods**) and those that pass this additional test (p < 0.05, red dots) are labeled. *Right*, normalized expression levels.

We next sought to find genes whose expression levels in starter cells associated with the number of inferred presynaptic partner cells (Fig. 3j). We hypothesized that differences in the expression of genes promoting or restricting synaptogenesis or dendrite growth influence the number of presynaptic cells innervating each starter cell. We focused our comparisons at the most granular cell subtype level and narrowed our testing to those genes sufficiently expressed and skewed in aggregate across presynaptic network size groups (**Methods**). We used permutation to create negative-control distributions in which each starter cell RNA profile was replaced by a randomly selected presynaptic RNA profile of the same type.

Across eight starter cell subtypes, 13 genes exhibited differential expression across presynaptic network size categories (p < 0.05, Wilcoxon Test; Fig. 4c; Extended Data Fig. 8b,c). Though this did not exceed the number of genes nominated in permuted analyses (mean±sem = 15.6±1.3), independent biological evidence strongly supported roles for two of the most strongly differentially expressed genes, both of which were more highly expressed in starter cells with large networks (relative to cells with small networks) and have described roles in promoting dendritic growth or synapse formation through developmental loss-of-function or ectopic overexpression experiments. *Arpp21* – which was upregulated ∼16 fold in postsynaptic *Pvalb*+ interneurons with large presynaptic networks – encodes an RNA binding protein that promotes dendritic growth by activating translation of target RNAs and whose cell-to-cell dynamic range might be extended due to intronic-encoded inhibitory microRNA^23^. *Cdh13* – which was upregulated ∼15 fold in postsynaptic*Tfap2d*+ glutamatergic neurons with large presynaptic networks – encodes an atypical protocadherin, one of four genes previously identified as driving synaptogenesis in a large-scale neuronal RNAi screen^12^.

### RNAs correlated with rabies virus transmission implicate synaptic function

Accurate interpretation of how rabies-inferred synaptic networks relate to actual synaptic connectivity and function is critically limited by our incomplete understanding of the molecules and cellular processes through which rabies enters, exits, and interacts with diverse host brain cell types. To determine which RNAs and biological pathways contribute to rabies transmission, we leveraged asynchronous development and variable rabies transmission in cultured cells to identify gene expression patterns that correlated with increased infectivity along the developmental trajectory stretching from neural precursor cells into mature SPNs (Fig. 5a,b). We strictly ordered each of the 32,503 scRNA profiles in pseudotime using Monocle3^24^ and confirmed the expected developmental processes through Gene Ontology Biological Pathway (GOBP)^25, 26^ enrichment analysis of co-regulated genes (∼25% of the coding genome; n= 7,844 genes; Extended Data Fig. 9a).

**Figure 5.**
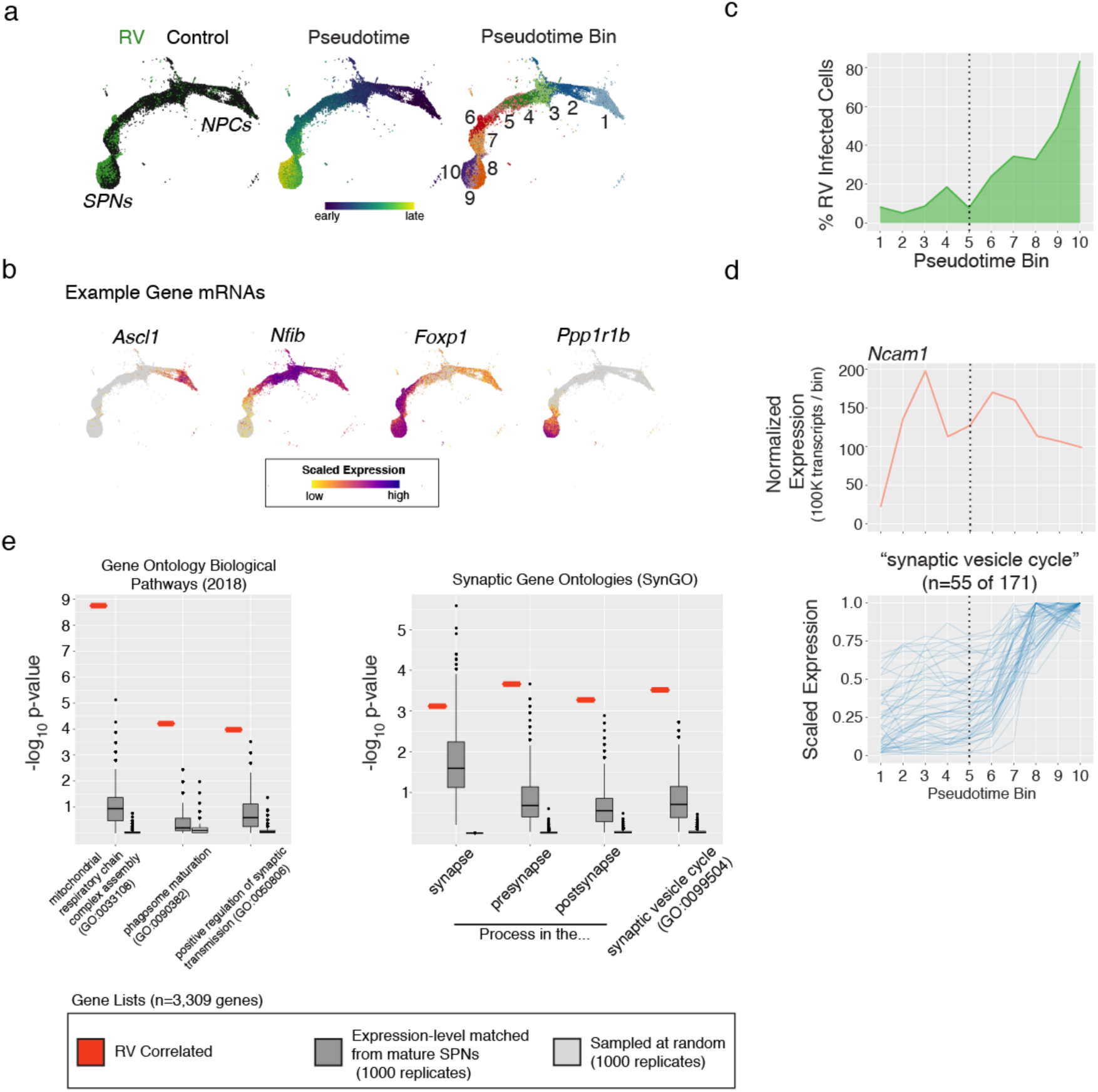
The developmental emergence of rabies transmission co-occurs with the maturation of synaptic function. **a.** UMAP embedding of scRNA profiles (n=32,503) along a trajectory of development from immature neural precursor cells (NPCs) to mature spiny projection neurons (SPNs). *Left*, color-coded by rabies virus infected (ie SBARRO; n=8,837 profiles) or uninfected control cells from paired cultures (n= 23,666 RNA profiles); *Middle*, pseudotime; *Right*, pseudotime bins (n=10). **b.** Example expression plots for four developmentally regulated genes. **c.** For each pseudotime bin, the percentage of scRNA profiles corresponding to rabies virus infected SBARRO cells over the total number of all cells. **d.** RNA levels across pseudotime bins. Of four described rabies receptors ^8^, *Ncam1* (*top*) has the only appreciable expression. *Ncam1* expression precedes the major developmental increase in rabies transmission. The subset of genes (n=55 of 171) in the “synaptic vesicle” SynGO category (*bottom*) whose RNA levels correlate with rabies virus infectivity (n=3,309 genes total). **e.** Gene Ontology Biological Pathways (GOBP) and Synaptic Gene Ontology (SynGO) analyses for rabies-infectivity correlated genes (n=3,309; **Methods**). *Left*, analyses were conducted on the correlated gene list (red) as well as two sets of control genes (each with 1,000 replicates of n=3,309 genes). In the “Expression-matched” set (dark grey), genes were selected from the mature SPN metacell (pseudotime bin = 10) in a manner that matched expression levels of the correlated genes. In the “Random” set (light grey), genes were selected at random from those for which expressed RNA was detected. P-value distributions for n=3 GO- BP categories (*middle*) and n=4 SynGO categories (*right*) for which the rabies- infectivity correlated gene set was statistically enriched.

We identified n=3,309 genes with RNA levels that correlated (r > 0.75) with increased rabies transmission (Fig. 5d,e and **Methods**). Interestingly, *Ncam1* mRNA was the only one of four described rabies receptors^8, 27, 28^ with appreciable expression in these experiments, and appeared in cells before high rates of infectivity, suggesting NCAM1 protein alone is not sufficient for rabies transmission (Fig. 5d). To discover which cellular processes might endow infectivity, we performed GOBP with the gene set we identified and compared the results to control gene sets sampled at random or from expression- matched mature SPN profiles. We identified selective enrichments in 1) “mitochondrial respiratory chain complex assembly”; 2) “phagosome maturation”; and 3) “positive regulation of synaptic transmission”, which were absent from control gene sets (Fig. 5d). To refine which synaptic processes were implicated in infectivity, we queried synaptic gene ontologies^29^ and identified selective enrichment for “synaptic vesicle cycle” (for which n=55 of 171 genes were correlated with infectivity). This analysis suggests that, in addition to the expression of viral entry receptors, operational synaptic transmission is critical for inter-cell rabies transmission and nominates specific genes implicated in the onset of synaptic transmission and rabies entry (Extended Data Fig. 9b).

## DISCUSSION

Our understanding of how synaptic networks emerge during development and how they are regulated by genetic and biological programs will benefit from measurements of synaptic connections that are systematic, quantitative, and connected to detailed molecular profiles of individual cells. Comprehensive characterization of the synaptic organization of neural circuits is challenging with current electrophysiological and anatomical methods, due to the small sizes of synapses, expansive geometries of axons and dendrites, and lack of knowledge of the cell subtypes involved. Such limitations have tended to separate synaptic-network biology from other subfields of neuroscience that are adopting highly parallel approaches for characterizing molecular repertoires^30–33^ or neural activity in many individual cells^34^.

Here we demonstrate that synaptic networks can be reconstructed from scRNA-seq data, thus allowing direct connectivity relationships to be inferred across thousands of individual cells for which genome-wide RNA expression has also been ascertained. Our data suggest that, during synaptogenesis *in vitro*, connectivity is shaped by cell type in a quantitative rather than qualitative way. Individual starter cells had considerable variance in their number of presynaptic partners, which appeared unrelated to the degree of infection or innate immune response, but partially explained by neuron type and gene expression patterns. We found that *Arpp21* and *Cdh13* had higher expression within starter cells with more presynaptic partners. Interestingly, *Cdh13* – an atypical transmembrane protein of the Cadherin superfamily – was previously identified as a key postsynaptic gene driving both excitatory and inhibitory synaptogenesis through a systematic RNAi screen^12^. In addition, *Arpp21* overexpression or knock-out bidirectionally controls the size and complexity of pyramidal neuron dendrites during postnatal development, likely by potentiating the translation of bound mRNA species that promote dendritogenesis^23^. These proof-of-concept observations suggest that extant molecular heterogeneity may associate with different properties of a given cell’s presynaptic network and that SBARRO analyses are a means to access and quantify such relationships.

We designed SBARRO to be adaptable to emerging single-cell genomic technologies. For example, methods enabling single-cell spatial transcriptomics^35–37^ or *in situ* sequencing^38, 39^ will allow the locations and anatomical properties of SBARRO cells to be mapped *in vivo* without cell loss. Moreover, long-read RNA isoform sequencing^40, 41^ could address long-standing hypotheses for how alternative splicing helps generate an extracellular adhesion code between synaptically connected cells within and across cell types^42^.

Unknown features of rabies cell biology represent a current limitation in the interpretation of SBARRO datasets. A detailed understanding of how our inferred, digital monosynaptic relationships relate to extraordinarily diverse and highly dynamic synaptic structures requires a comprehensive description of how rabies interacts with and transits between host brain cells of different types. On one hand, previous studies provide direct and circumstantial evidence that suggest, at least to a first approximation, that rabies transmission events are selective for synapses made directly onto infected neurons^43, 44^. Among postsynaptic neurons of the same class with spatially intermixed dendrites, presynaptic labeling respects synapse-selective motor arcs in the spinal cord. Similarly, in primary visual cortex, intermingled layer 2/3 glutamatergic neurons distinguished by firing properties to visual cues, appear to inherit those selective properties from presynaptic cells labeled by rabies infection^45, 46^. On the other hand, the efficiency of rabies transmission can be very low for certain cell-type-specific axons^47^ and appears to be modulated by presynaptic firing rate^48, 49^. Our correlative molecular data suggest developmentally mature presynaptic function is critical for rabies uptake in neurons (Fig. 5). However, the extent to which rabies egress and entry exclusively use synapse-associated processes; occur through direct synaptic contacts; and are affected by neural activity across diverse brain circuits, all remain to be firmly established^48^. Moreover, it will be necessary to study the ways in which infection alters host cells’ molecular programs, as these alterations could affect synapse-associated processes.

Rabies infection of non-neuronal cell classes, such as astrocytes and polydendrocytes, is a minor yet clear feature of our *in vitro* experiments and is also observed *in vivo* ^10, 52, 53^. While both cell types interact intimately with synapses, especially during development, more experiments are necessary to understand the molecular mechanisms underlying rabies transmission across non-neuronal classes. Single-cell, single-virion inferences of these interactions may offer valuable insight: RABID-seq^6^ analyses suggest rabies can be transmitted from infected astrocytes into presumed physically-adjacent microglia and that specialized host cell signatures are associated and detectable with these interactions. What role synapses play in these glial interactions is not clear. (Microglia were not present in our experiments *in vitro*).

Mammalian synaptogenesis is particularly challenging to study with traditional methods due to the many cell types and molecules involved, its protracted nature in space and time, and intrinsic noise that arises from being a competitive, cell-to-cell process. By facilitating connectivity inferences and RNA sampling from the same individual cells, we hope that fast, scalable, all-molecular approaches such as SBARRO – which may be eventually deployed in non-destructive ways^50^ – can complement established connectomic technologies based on super-resolution imaging of synaptic anatomy.

## DATA AVAILABILITY

The sequencing data reported in this paper are in the process of being uploaded to GEO. A GEO accession number will be provided upon completion.

## CODE AVAILABILITY

Software and core computational analysis to align and process scRNA-seq reads are freely available: https://github.com/broadinstitute/Drop-seq/releases. Other custom code available by request.

## ACKNOWLEDGMENTS

This work was supported by the Broad Institute’s Stanley Center for Psychiatric Research and a Helen Hay Whitney Postdoctoral Fellowship to AS. The authors thank Frank Koopmans for SynGO analysis scripts, Dr. Fenna Krienen, Dr. Marta Florio, Dr. Christina Usher and other members of the McCaroll lab for helpful advice.

## AUTHOR CONTRIBUTIONS

AS conceived the idea, designed and supervised experiments, analyzed the data and drafted the manuscript. Plasmid barcoding, CV and AS. Rabies virus packaging, KWH, AP, CV, AS and BLS. Cell culture, KS and CS. scRNA-seq experiments/analysis and molecular protocols, CS, KS, HS and AS. Algorithm development, SAM, JN and AS. Analysis software, SK. Computational support, AW. Manuscript preparation, AS and SAM with input from other authors.

## AUTHOR INFORMATION

Correspondence and request for materials should be addressed to AS or SAM.

## CORRESPONDENCE

Arpiar Saunders; Steve McCarroll.

## METHODS

### Barcoding rabies virus plasmids and RNA genomes

Rabies virus rescue encapsidates RNA genomes from DNA templates^51^. Generating rabies virus particle libraries with millions of unique and similarly abundant genomic barcodes presents a two-part challenge not encountered when rescuing a single genomic species. First, a plasmid library carrying hyper-diverse barcoded DNA genomes is created *ab initio*. Second, barcode loss and abundance skews must be minimized during plasmid amplification (in bacteria) then in rabies virus rescue and replication (in mammalian cell culture). To address these challenges, custom protocols were developed to 1) introduce barcode sequences into DNA plasmids using PCR (achieving near-theoretical levels of plasmid-to-plasmid barcode diversity; Fig. 1f and Extended Data Fig. 2a); 2) more uniformly amplify plasmid DNA through optimized bacterial transformation and plate-based growth conditions (Extended Data Fig. 2c); and 3) rescue rabies virus (with native or pseudotyped coat proteins) in ways that mitigate distortions in barcode representation, initially created by the very low-probability of individual rescue events^10^ and then exacerbated by biases in viral replication. Our 7-9 day protocol is three-fold faster and achieves titers equivalent or higher than published protocols (1×10^8-9^ IU/mL; Extended Data Fig. 2d)^15, 52^. Details for each of the three protocol steps are found in the sections below. Barcodes present in DNA plasmids and RNA genomes were quantified through sequencing-based approaches in which oligonucleotide probes containing unique molecular identifier (UMI) sequences were hybridized to barcode-adjacent sequences and then polymerase-extended through the barcode region (Extended Data Fig. 1c and **Supplementary Table 1**); the resulting paired UMI-barcode sequences were used to count individual molecules. Inflation of barcode sequences and UMI counts due to mutations arising during library amplification and Illumina sequencing were accounted for (see description in Results) using a custom algorithm for post-hoc mutation correction. See “*Quantifying barcodes from plasmids and rabies virus genome”* section below for details.

#### PCR-based plasmid barcoding

To generate plasmid libraries in which individual circular plasmids encode unique barcode sequences, we developed a PCR-based molecular workflow in which a bipartite barcode cassette can be targeted to arbitrary regions of a non-barcoded plasmid template (Extended Data Fig. 1b). We applied our system to the SAD-B19 genome plasmid in which the G gene has been replaced by *EGFP* (cSPBN-4GFP, Addgene #52487^15^), targeting the barcode cassette to the 3’ UTR of *EGFP* adjacent to the viral polyadenylation sequence^53^. To introduce each half of the barcode cassette, whole-plasmid PCR was performed with forward and reverse primers targeting the desired region. Each primer contains 3’ plasmid-complementary sequence followed by a 5’ tail with 10 bps of random nucleotides further flanked by a restriction cassette which includes the PlutI (“GGCGCC”) restriction site (pSPBN-GFP Barcoding, Forward Primer: B19_barcode_F, Reverse Primer: B19_barcode_R; **Supplementary Table 1**). During PCR (See *“Barcoding PCR”* protocol), each round of primer hybridization and extension introduces a unique barcode, resulting in a linear, double-stranded amplicon collection in which unique 10 bp barcodes have been introduced into the 5’ terminus of each DNA strand. The desired ∼14.5 kb amplicons were size-selected using standard low-gel agarose (Sigma-Aldrich, A9414) electrophoresis and cleaned (Zymo Research, Gel DNA Recovery Kit #D4001), then re-cleaned and concentrated to >200 ng/μl (Zymo Research, DNA Clean & Concentrator-25 #D4033). To efficiently circularize the amplicons using the barcode restriction cassette and to remove remaining template plasmid and unwanted linear products, we developed a series of enzymatic reactions that consecutively performed in the same tube, saving time and avoiding DNA damage and loss due to repeated purification (See *“Plasmid Circularization Protocol”*). Briefly, DpnI digest removes remaining methylated plasmid DNA; PlutI restriction and T4 ligation circularize the amplicons thus covalently bonding each of two 10 bp barcodes into a 36 bp barcode cassette; and RecBCD selectively degrades linear DNA over circularized plasmid containing non-complementary barcode sequences, typically enriching the percentage of circularized product ∼3.5 fold (from ∼20±0.7% to 71±4%, n=6 experiments, ± denotes s.e.m).

##### Barcoding PCR

1. PCR (25 ml reaction)

1. PCR (25 μl reaction)
| Reagent | Volume (μl) |
| --- | --- |
| Q5 High-Fidelity 2X Master Mix (NEB, #M0515) | 12.5 |
| Forward primer (B19_barcode_F) | 1.25 |
| Reverse primer (B19_barcode_R) | 1.25 |
| Template plasmid (cSPBN-4GFP, 0.4 ng/μl) | 5 |
| Water | 5 |

##### Plasmid Circularization Protocol

1. Digest (50 μl reaction)

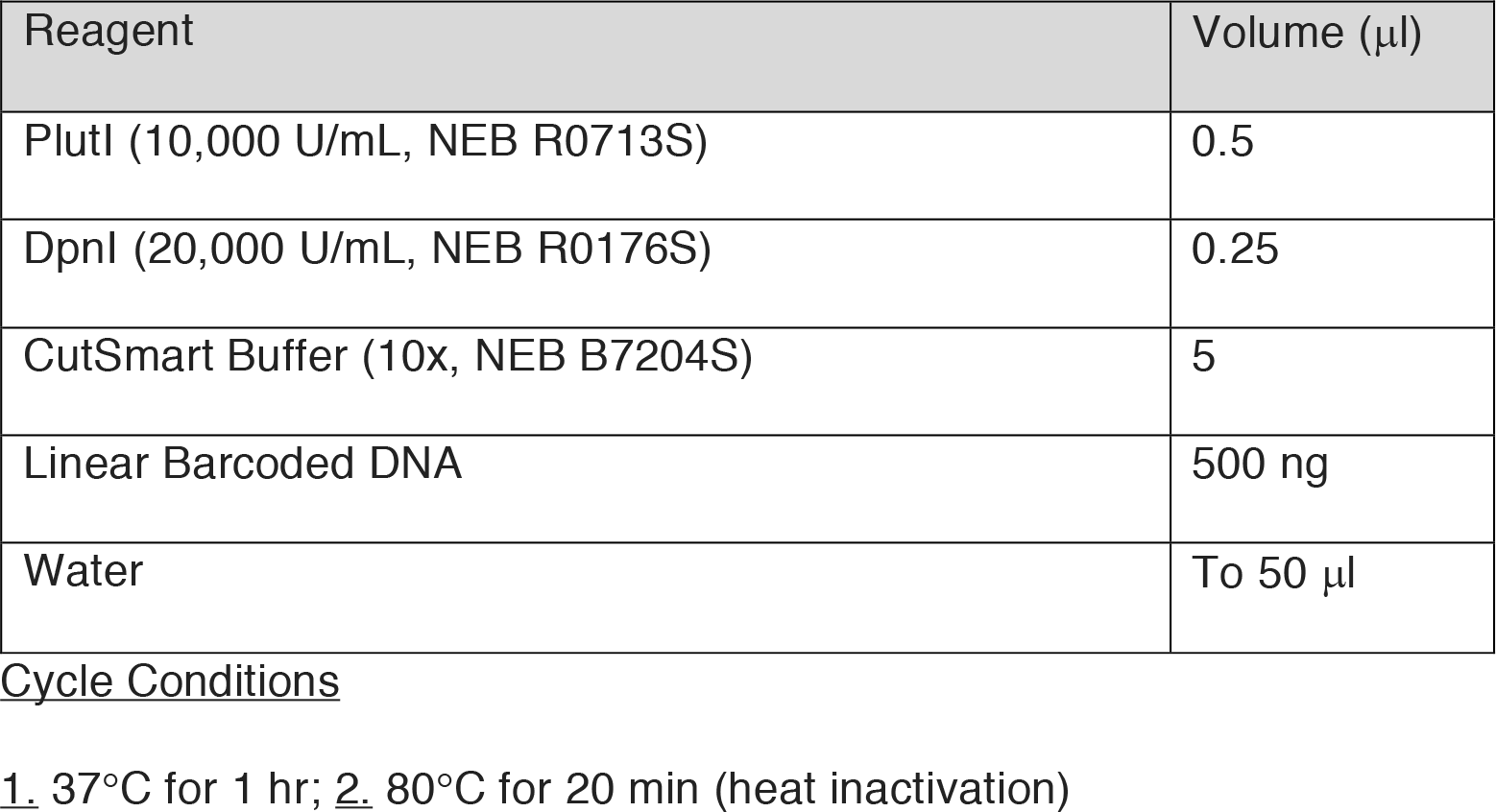

| Reagent | Volume (μl) |
| --- | --- |
| PlutI (10,000 U/mL, NEB R0713S) | 0.5 |
| DpnI (20,000 U/mL, NEB R0176S) | 0.25 |
| CutSmart Buffer (10x, NEB B7204S) | 5 |
| Linear Barcoded DNA | 500 ng |
| Water | To 50 μl |

Cycle Conditions
1. 37°C for 1 hr; 2. 80°C for 20 min (heat inactivation)

2. Ligation (spike-in, +5.1 ml)

| Reagent | Volume (μl) |
| --- | --- |
| T4 DNA Ligase (10,000 U/ml, NEB M0202T) | 0.1 (200 U) |
| ATP (10 mM, NEB P0756L) | 5 |

Cycle Conditions
1. 4°C for 2 hrs; 2. 65°C for 20 min (heat inactivation)

3. Circular plasmid enrichment (spike-in, +8.8 ml)

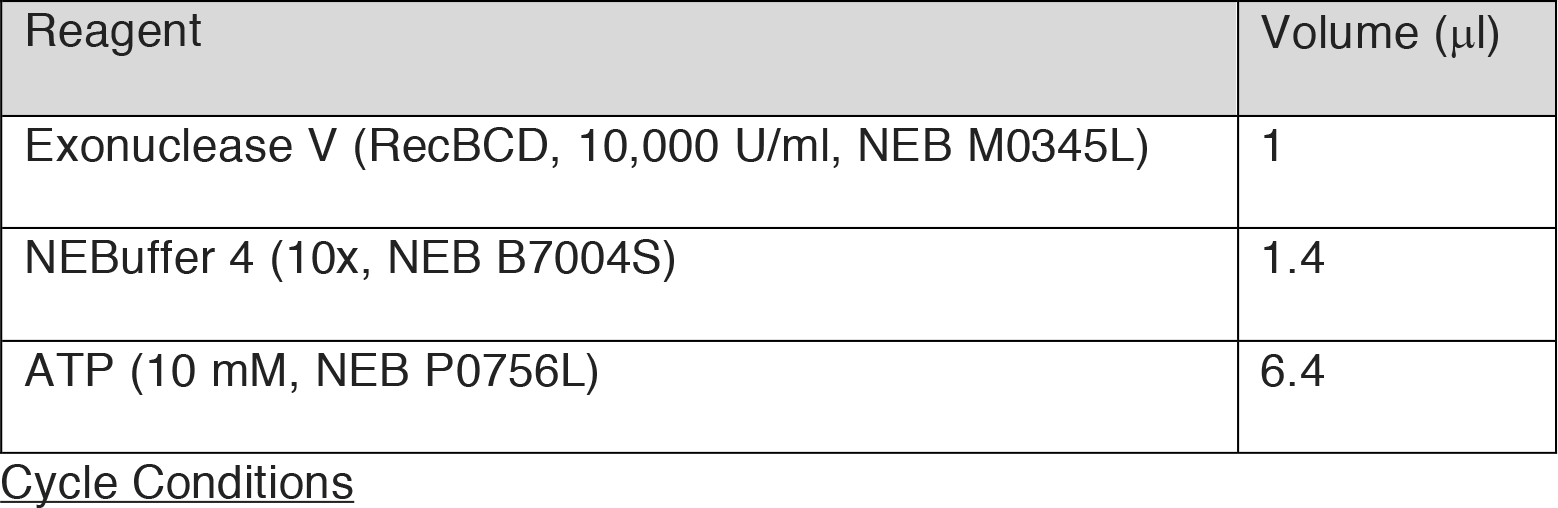
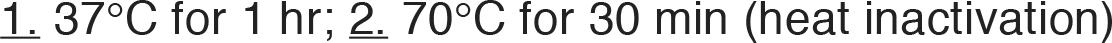

| Reagent | Volume (μl) |
| --- | --- |
| Exonuclease V (RecBCD, 10,000 U/ml, NEB M0345L) | 1 |
| NEBuffer 4 (10x, NEB B7004S) | 1.4 |
| ATP (10 mM, NEB P0756L) | 6.4 |

Cycle Conditions
1. 37°C for 1 hr; 2. 70°C for 30 min (heat inactivation)

#### Amplifying barcoded DNA plasmids

Rabies virus rescue requires tens of micrograms of supercoiled rabies virus genome plasmid for cell transfection. To amplify and supercoil DNA plasmid libraries carrying hyper-diverse barcodes in a manner that minimizes loss and skew of barcoded plasmid representation, we transformed eight vials of chemically competent One Shot OmniMAX 2 T1^R^ cells (ThermoFisher Scientific, C854003) in parallel each with 200 ng of DNA from the *Plasmid Circularization Protocol*. After 1 hour of recovery growth, cultures were combined and 2 ml of the cell mixture was spread over n=8 large plates (24.5 x 24.5cm; Corning, CLS431111^54^) containing LB Agar (Sigma-Aldrich, L2897) and Ampicillin (100ug/mL; Sigma-Aldrich, A5354) and grown over night at 37°C. Colonies were scraped from each plate with 15 ml LB and pelleted through centrifugation (6000g for 15 min at 4°C). Plasmids were isolated from cell pellets using the EndoFree Plasmid Maxi Kit (0.45 g cells/column; Qiagen, 12362). Sequencing-based barcode quantification (see below *Quantifying barcodes from plasmids and rabies virus genomes*) comparing plasmids prepared from pooled transformants grown on plates (as described) versus liquid culture (250 ml) demonstrated that plate-based growth dramatically reduced overrepresentation of plasmid barcodes (Extended Data Fig. 2c), presumably by homogenizing clonal growth rates.

#### Rescuing barcoded rabies virus libraries

*De novo* rescue of negative-stranded RNA viruses requires transfection-based encapsidation of positive-stranded RNA genomes with N, P and L proteins; the minimal replication-competent nucleocapsid ^51^; Genomes lacking the G gene additionally require G protein such that replicating particles can spread cell-to-cell ^15^. Two properties of rescue create challenges for generating particle libraries with millions of unique and uniformly abundant genomes. First, cells in which rescue events occur are rare (<1:10,000 transfected cells ^10^), creating a limited number of cellular environments in which encapsidation can occur. Second, state-of-the-art packaging protocols serially infect fresh cultured cells to increase viral titer; increasing, with each passage, the opportunities for individual clones gain a replication advantage. To develop a rabies virus rescue protocol for barcoded genomes, we first systematically characterized how barcode abundances behaved after transfection and each passage stage of a widely used protocol^15^. We observed that 1) minimally and on average, hundreds of unique rabies virus genomes were encapsidated per encapsidation-competent cell and that 2) viral passages tended to reduce the number of unique barcoded genomes and distort their relative abundances (Extended Data Fig. 3). Therefore, we increased the total number of encapsidation- competent cells by optimizing large-scale rabies virus transfection and created a one- step rescue protocol capable of generating rabies virus libraries with millions of unique genomes at similarly high titers (∼2.5 x 10^9^ IU/ml) but 4-fold faster (5 days) than published protocols ^15, 52^. Pseudotyping with non-native coat proteins requires an additional 6 days. Specifically, poly-L-lysine (Sigma-Aldrich, P4707) coated T-225 flasks containing 85-95% confluent HEK-293T/17 cells (ATCC, CRL-11268) were each transfected (Xfect, Takara #631318) with a DNA cocktail containing the 1) the barcoded rabies virus plasmid library (131.36 μg) and CAG-promoter driven plasmids for T7 polymerase (23.66 mg, Addgene 59926) and SAD-B19 helper proteins (N, 52.11 mg, Addgene 59924; P, 30.15 μg, Addgene 59925; L, 23.70 mg, Addgene 59922; G, 20.26 μg, Addgene 59921). Cells were maintained with DMEM with GlutaMAX supplement, pyruvate, high glucose media (Thermo Fisher Scientific, 10569010) supplemented with 5% fetal bovine serum (Thermo Fisher Scientific, 10082147) and 1x antibiotic-antimycotic (Thermo Fisher Scientific, 15240062) and incubated at 35°C with 5% CO2. Five days post-transfection, culture media was collected for either 1) unpseudotyped rabies virus recovery or 2) EnvA pseudotyping. For EnvA pseudotyping, BHK-EnvA cells (Columbia Univ. Zuckerman Virus Core), initially grown in 15 cm dishes (Corning, 08-772-24) to 85-95% confluence, were infected with filtered media (0.22 µm PES; Corning, 431097) from the transfected T-225 plates; cells are then rinsed, pelleted and re-plated first in 15 cm plates then again in T-225 flasks. Specifically, following 6 hours of incubation with particle-containing media, cells from each plate are rinsed with two rounds of cold DPBS (+Ca, +Mg), trypsinized with 5 mL of trypsin-EDTA (Thermo Fisher Scientific, 25300-054) for 30 seconds at 35°C, pelleted with centrifugation (300 *g*, 4 min) in DMEM + 10% FBS and then re-plated in 15 cm plates and allowed to incubate overnight (∼16-24 hours) before being re-plated in T-225 flasks with DMEM + 5% FBS. T-225 plate media is supplemented with 3-5 mL of DMEM + 5% FBS each day for 4 days before being collected and concentrated. Specifically, collected media is incubated with benzonase nuclease (1:1000 dilution; Millipore Sigma, 70664) for 30 minutes at 37°C and filtered (0.22 µm PES). For ultracentrifugation (Beckman Coulter, SW32Ti rotor), 2 mL of 20% (w/v) sucrose in DPBS (-Ca, -Mg) is prepared in ultracentrifuge tubes (Beckman Coulter, 344058) to which the divided EnvA- pseudotyped viral media is added before pelleting (20,000 RPM for 2 hours at 4°C). Residual media is removed and viral pellets are each resuspended in 15 µL of DPBS (-Ca, -Mg) on ice before orbital shaking at 4°C for 8 hours. Volumes are then combined, aliquoted, and stored at -80°C. Titers were established by quantifying infected HEK-TVA and HEK-293T/17 cells in 12-well plates (80% confluence) using serial dilutions. To ensure EnvA-pseudotyping was complete, < 2 HEK-293T/17 cells per well were tolerated following infection with 1 µL with full-strength sample.

#### Quantifying barcodes from plasmids and rabies virus genomes

UMI-based counting of barcodes from DNA plasmids and RNA genomes was accomplished with similar molecular (See “*UMI-based counting of genome and plasmid barcodes*” and Extended Data Fig. 2) and informatic workflows (See “*UMI-based counting of genome and plasmid barcodes*”). RNA genomes were extracted using the ZR Viral RNA kit (Zymo Research, R1041) from particles ascertained from 1) end stage high- titer viral aliquots or 2) from cell culture media used for rabies virus rescue after PEG- based precipitation (Abcam, ab102538) and quantified using the High Sensitivity RNA ScreenTape assay (Agilent, 5067-5579). To count barcode abundances of individual RNA genomes or DNA plasmids, an oligonucleotide (B19_UMI_F) containing a SMRT PCR handle, 12 bp UMI and 33 bps of barcode-adjacent homologous sequence were hybridized then polymerase-extended through the barcode region. Remaining RNA genomes were selectively digested using RNase H (New England Biolabs, M0297S) and reactions were cleaned with Agencourt AMPure XP beads (1:1 volume; Beckman Coulter, A63881) retaining first-strand cDNA. The UMI-tagged genomic cDNA or plasmid DNA strands were then selectively amplified (14-18 PCR cycles; 16 median) using primers which introduce the Illumina P5 (P5-TSO_Hybrid) and indexed P7 (P7i1-L5UTR_seq) sequences. Amplicon libraries were sequenced on an Illumina MiSeq or NextSeq550 using a custom primer (Read1CustomSeqB) to seed 110 Read 1 cycles. Base pairs (bp) 1-12 were assigned as the UMI. The two 10 bp viral barcodes were informatically extracted from the barcode cassette using a custom algorithms based on local sequence alignment algorithm and (“*TagReadWithRabiesBarcodes*” & “*FilterValidRabiesBarcodes*”). To account for artifactual barcode sequences created by mutations acquired during the library amplification and sequencing, we developed an algorithm to identify and collapse “families” of barcodes with similar sequences likely related through acquired mutations *(“CollapseTagWithContext, MUTATIONAL_COLLAPSE=true”*). Specifically, after ordering barcodes most to least abundant, we considered each barcode as a “parent” and identified “siblings” sequences within Hamming distance of 1 of the “parent” barcode. The process was then iterated for each new “sibling” until no new “siblings” were discovered. The entire barcode family was assigned the sequence of the “parent.” UMI-parent barcode sequence pairs were then used to count each “parent” barcode in the library (after collapsing UMI-barcode sequences in which the UMIs associated with the same “parent” were Hamming distance <= 1). This approach drastically reduced the inflation of barcode sequences and counts due to library preparation and sequencing (Extended Data Fig. 2b).

##### UMI-based counting of RNA genome and DNA plasmid barcodes

1a. RNA Genomes - UMI Hybridization (24 μl reaction)

1a. RNA Genomes - UMI Hybridization (24 μl reaction)
| Reagent | Volume (μl) |
| --- | --- |
| Rabies ssRNA genomes or dsDNA plasmid (5-25 ng) | -- |
| UMI oligo (10 μM; B19_UMI_F) | 4 |
| dNTPs (10 mM; ) | 4 |
| Water | To 24 µl |

Cycle Conditions
1. 72°C for 4 min; 2. 4°C

1b. RNA Genomes - Reverse Transcription (spike-in, +16 ml)

| Reagent | Volume (µl) |
| --- | --- |
| 20% Ficoll PM 400 (Sigma Aldrich, 26873-85-8) | 4 |
| NxGen RNase Inhibitor (Lucigen, 30281-2) | 0.5 |
| Maxima H- RT (Thermo Scientific, EP0751) | 2 |
| 5x Maximal H- RT Buffer | 8 |
| Water | 1.5 |

Cycle Conditions
1. 42°C for 90 min; 2. 85°C 5 min (inactivation)

1c. RNA Genomes – RNase H Treatment (spike-in, +2 ml)

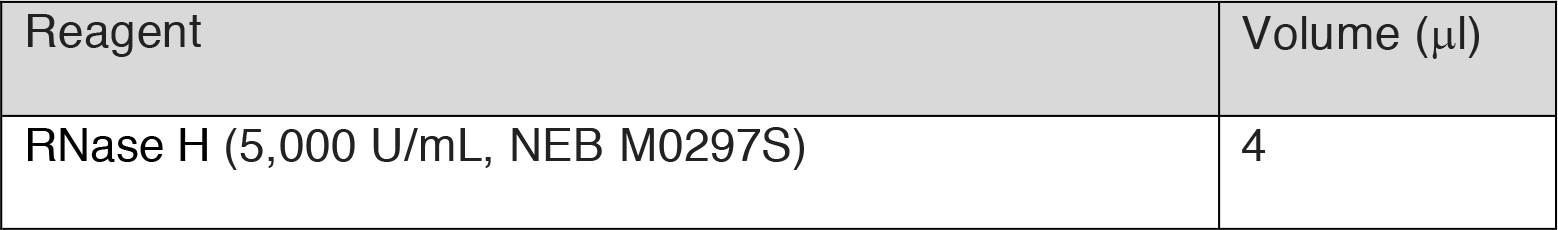

1. DNA Plasmids - UMI Hybridization & polymerization (50 μl reaction)

| Reagent | Volume (µl) |
| --- | --- |
| dsDNA plasmid (5-25 ng) | -- |
| 2x KAPA HiFi HotStart ReadyMix<br>(Kapa Biosystems, KK2602) | 25 |
| UMI oligo (10 µM; B19_UMI_F) | 2 |
| Water | To 50 µl |

2. Illumina Adaptor PCR (50 μl reaction)

| Reagent | Volume (µl) |
| --- | --- |
| UMI-tagged DNA | 5 |
| P5 Primer (10 µM; P5-TSO_Hybrid) | 1 |
| P7 Primer (10 µM; P7i1-L5UTR_seq) | 1 |
| 2x KAPA HiFi HotStart ReadyMix<br>(Kapa Biosystems, KK2602) | 25 |
| Water | To 50 µl |

Cycle Conditions
1. 98°C for 30s; 2. 98°C for 10s; 3. 72°C for 30s; 4. 72°C for 60s; 5. Go to Step 2-4,
13-17 times; 6. 72°C for 120s

### Synaptic cell culture

Cells were dissociated from the cortex or striatum of embryonic day 16 (E16) C57Blk6/N mouse brains and maintained for 14 days *in vitro* (DIV). rAAVs were transduced on DIV 5 to functionalize “starter” cells. On DIV 12, EnvA-RV*dG*-*EGFP*VBC libraries were transduced and infection was allowed to proceed for 72-96 hours before scRNA-seq libraries were generated. Pregnant C57Blk6/N dams (Charles River Laboratories) were heavily anesthetized by isoflurane inhalation, decapitated and the brains of embryonic pups (litter size = 4-9) removed in ice-cold 1X Dissociation Media (“DM”; containing (in mM): 10.52 MgCl2 (Sigma-Aldrich, M2393); 10.53 HEPES (Sigma-Aldrich, H3375); 1.32 Kynurenic Acid (Sigma-Aldrich, K3375) in HBSS (Thermo Fisher Scientific, 14175079)) in which cortex or cortex and striatum were dissected from each brain, pooled and incubated in sterile-filtered (0.22 μm; Corning, 431097) DM+Papain/L-Cysteine (3.4 units Papain and 0.172 mM L-Cysteine; Worthington Biochemical, LK003178) for 3-5 min at 37° C. Brain tissue is then washed twice with 2-3 mL sterile-filtered DM+Trypsin Inhibitor (1 mg/mL; Sigma Aldrich, T9253) and incubated in the 3^rd^ wash for 3-5 min at 37° C. DM+Trypsin Inhibitor is replaced with 6 mL of sterile-filtered “Plating Media” (“PM”; containing: DMEM (ATCC, 30-2002) and 10% FBS (ATCC, 30-2020)) in which digested brain volumes are titrated into single cells with a pipetteman equipped with a 5 mL pipet tip. Cell concentration was measured by diluting cells 1 to 5 in PM, mixed with an equal volume of 0.4% Trypan Blue (Thermo Fisher Scientific, 15250-061), and quantified using the Countess II Automated Cell Counter (Life Technologies). Each well of 6-well cell culture treated plates coated with 0.1% Poly-L-ornithine (3 ug/mL; Sigma Aldrich, P4957) were seeded with ∼750K cells and maintained with sterile-filtered neurobasal medium (Thermo Fisher Scientific, 21103049), supplemented with serum-free B-27 (Thermo Fisher Scientific,17504044), GlutaMAX (Invitrogen, 35050061) and Penicillin:Streptomycin (VWR, 45000-652). For imaging experiments, cells were seeded on glass coverslips (Fisher Scientific, 12-546) coated with 0.1% Poly-L- ornithine and Laminin (5 ug/mL; Thermo Fisher Scientific, 23017015). On DIV 5, a cocktail of three rAAVs was used to functionalize starter cells. Our starter cell strategy was designed to deliver consistent, high MOIs of rAAV per starter cell while flexibly controlling the number of starter cells in each culture through Cre delivery. Specifically, 1 μl CAG-Flex-TVA-mCherry (“TCB”; serotype, 2-9; titer, 2.2×10^13^ genomes/mL; MOI, ∼2.9e^4^; UNC Vector Core) and 1 μl CAG-Flex-B19G (serotype, 2- 9; titer, 1.6×10^13^ genomes/mL; MOI, ∼2.1e^4^; UNC Vector Core) were added to each well along with 1 μl of Syn1-EBFP-Cre (serotype, 2-1; titer, 6×10^12^ genomes/mL; MOI, ∼8-0.08; Addgene, 51507-AAV1) delivered at full strength or diluted 1:10^3^-10^4^. At DIV 12, 1 μl of EnvA-RV*dG*-*EGFP*VBC library (titer, 0.19-1.1×10^10^ IU/mL; Total cell MOI, 2.5- 14.8) was added to each well. Epifluorescence imaging was used to monitor the progress of infections, including starter cell locations and morphology (based on TVA- mCherry fluorescence) as wells as rabies virus transduction and spread from starter cells (based EGFP fluorescence). , EnvA-RV*dG*-*EGFP*VBC transduction was completely dependent on the TVA receptor, since no EGFP fluorescence was observed in equivalent experiments in which the CAG-Flex-TVA-mCherry rAAV was excluded. To prepare cultures grown on coverslips for imaging or *in situ* hybridization experiments, neurobasal medium was removed and each culture well was rinsed three times with 1X PBS and then fixed with fresh 4% paraformaldehyde at room temperature for 30 min, followed by three rinses with 1x PBS. For fluorescence imagining, coverslips were slide-mounted and nuclei counterstained using ProLong Gold Antifade (Thermo Fisher Scientific, P36934). For *in situ* hybridizations, cover slips were dehydrated through a series of brief (∼1 min) ethanol washes (50%, 70% and 100% EtOH) before being stored at -20°C in 100% EtOH.

### Sequencing single-cell mRNAs: host cell, rabies virus barcodes and recombined rAAV

scRNA-seq libraries were generated using the Chromium Single Cell 3’ v2 or v3 Chemistry platform (10x Genomics), prepared following kit guidelines, and sequenced to a depth of ∼45K reads per cell (Illumina NovaSeq 6000). Sequences were aligned using STAR v2.4.0a^55^ against a composite genome consisting of GRCm38.81, barcoded cSPBN-4GFP and rAAV accessory sequences (including the 3’ UTR and TVA-mCherry, rabies G and Cre coding sequences) using a workflow similar to that described for Drop-seq^56^. To create input cell suspensions, a protocol developed for the adult mouse brain^30^ was adapted for *in vitro* synaptic cultures. Culture wells were first incubated for ∼20 min at 37°C with 1.8 mL of Dissociation Media (DM) containing Papain and Protease 23 ^30^ until detachment of the cell monolayer. Cultures were gently swirled and incubated for an additional 5 min. Each well was then supplemented with 1 mL of DM before transfer into a 5 mL eppendorf tube in which cells were pelleted through centrifugation (300*g* for 5 min). The supernatant was removed and replaced with 1 mL of ice-cold DM in which cells were titrated by successively smaller bore polished glass Pasteur pipets. The cells were then re-pelleted and resuspended in 0.5 mL of DM before being filtered through a pre-wet 40 μm cell strainer (Corning, 352340). For SBARRO experiments, rabies infected cells were enriched from total cell suspensions through fluorescent activated cell sorting (FACS) using the MoFlo Astrios EQ cell sorter (Beckman Coulter; 70 mm nozzle) into 25 μl of DM. RV-derived EGFP fluorescence was used to gate for SBARRO cells. For experiments in which distinct scRNA-seq libraries were created for starter and presynaptic cells, mCherry fluorescence (driven by cre-recombined rAAV genomes encoding TVA-mCherry) was used as an additional gate to sort starter (GFP+/ TVA-mCherry+) or presynaptic (EGFP+/ TVA-mCherry-) populations. Post hoc FACS analysis was performed with FloJo software (BD Biosciences). For scRNA- seq libraries downstream of FACS, 1.7K-33K cells (based on FACS counts) were loaded per single-cell RNA capture reaction. To generate scRNA-seq libraries for which total cell suspensions were used as input, cell concentrations were quantified using the Countess II Automated Cell Counter (Life Technologies) and 10K-16K cells were loaded per single-cell RNA capture reaction. Rabies virus barcoded EGFP and cre-recombined TVA-mCherry rAAV 3’ mRNAs were independently amplified (See “*Selective mRNA Adaptor PCR*” protocol; Rabies EGFP: ∼255 bp amplicon, 10-14 cycles; rAAV TVA-mCherry: ∼1,125 bp amplicon, 34 cycles) from single-cell cDNA using primers that introduced the Illumina P5 site (P5-TSO_Hybrid) and indexed P7 site onto barcoded 3’ EGFP (BC_Seq_P7ìx’_GFP_v4c) or recombined TVA-mCherry (P7’x’_TCB_CreOn_v4). Barcoded EGFP and recombined TVA-mCherry libraries were multiplexed and sequenced separately on an Illumina NextSeq500 using a High Output 150 cycle Kit (Stock Read1 primer; Library concentrations: EGFP, 1.8 pM with 20% PhiX; TVA-mCherry, 0.4 pM with 50% PhiX; Cycle distributions: Read1=28, Read2=98, Index=8; Reads per library: EGFP, 43M-90M; TVA-mCherry, 121K-8.6M). To generate integer counts of recombined TVA-mCherry transcripts per cell, sequences generated from recombined TVA-mCherry library were aligned using STAR v2.4.0a ^55^ against a composite genome consisting of GRCm38.81, the RV*dG*- *EGFP*VBC genome and rAAV accessory sequences (including the 3’ UTR and TVA- mCherry, rabies G and Cre coding sequences). The sequences of UMIs associated with each gene and cell barcode were collapsed within an edit distance of 2. To quantify the number of TVA-mCherry mRNAs derived from cre-recombined rAAV genomes per cell, UMI counts mapping to the TVA-mCherry coding sequence or the 3’ UTR were summed. To discover and quantify the RV-derived VBCs in the 3’ UTR of EGFP mRNA, raw VBC sequences were informatically extracted from each read (as described above for plasmids and viral genome sequences). To accurately reconstruct and count VBCs in each single-cell, we leveraged the single-cell nature of the data to informatically account for two types of artifacts: 1) the inflation of barcode sequences generated by mutations during library amplification and sequencing and 2) swapping of non-adjacent VBC and cell barcode/UMI sequences due to strand displacement during PCR amplification (“CollapseTagWithContext, ADAPTIVE_EDIT_DISTANCE=true” & “BipartiteRabiesVirusCollapse”). To account for mutations, we assumed that in individual cells, closely related barcode sequences were likely to originate from mutations introduced during library preparation or sequencing rather than independent infections of rabies virus particles with similar 20 bp genomic barcodes. Thus we evaluated Hamming edit distance relationships across all sufficiently abundant VBCs (Inclusion Threshold: >= 3 (“No RG” experiments) or 5 (“SCC” Experiments) UMIs) found within each cell. From these edit distance distributions, many low-abundance “sibling” VBCs with sequences similar to a single, more numerous “parent” VBC were assigned the VBC of the “parent”; collapsing these mutationally-related VBC “families” corrected the strong artifactual correlation present in the raw data in which cells with more VBC UMIs also tended to have more unique VBCs (Extended Data Fig. 3a) reduced the number of included CBC-VBC counts by 82.4%. In the single-cell cDNA, cell barcode/UMI sequences are separated from the VBC cassette by >20 bps - including tracts of A/T homopolymers – providing an opportunity for mispairing of critical barcode sequences during PCR through strand- displacement or mispriming. To account for mispairing events, in cells with multiple VBCs, we developed a collapse algorithm based on fraction of shared UMI sequences (within edit distance 2) shared across each pair of VBCs. For pairs with >50% UMI sharing, the “sibling” VBC with fewer UMIs was assigned the VBC of the more abundant “parent”, enforcing that CBC-UMI barcodes should not be used by more than a single VBC. The ratio of within-cell VBC collapse events due to UMI sharing versus total CBC-VBC counts averaged 0.13±0.02 (s.e.m) across experiments; a correction which reduced the number CBC-VBCs counts by an additional 1%. Taken together, these two VBC collapse steps reduced included CBC-VBC counts by 83.4% as compared to the raw data - thus drastically altering the inferred groupings of single cells into networks – and also shaped within-cell VBC quantification, altering the UMI counts for ∼15% of VBCs (Change in VBC UMIs: mean, 5.7; median, 2).

Selective mRNA Adaptor PCR (50 μl reaction)

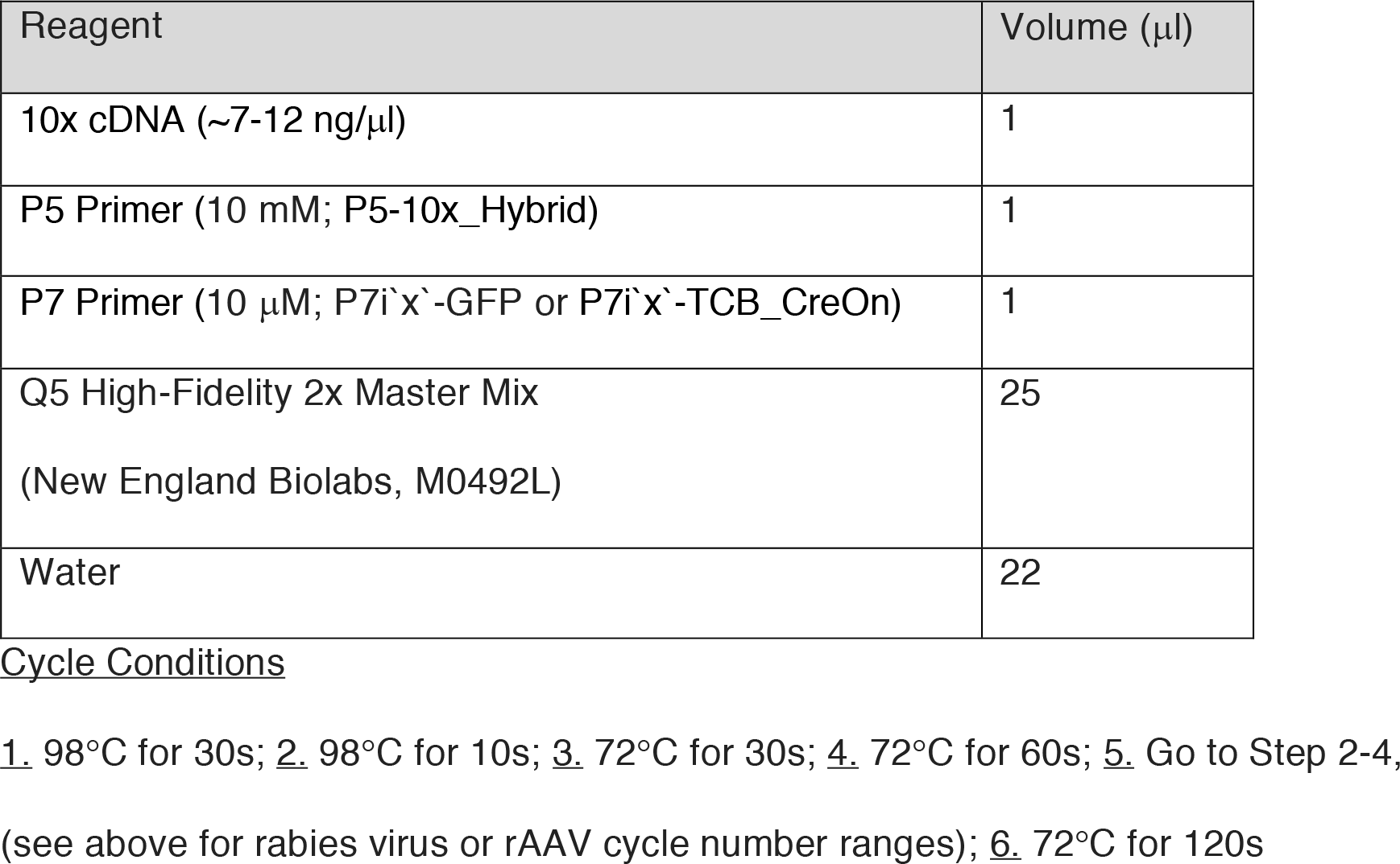

### Identification, clustering and analysis of host cell scRNA profiles

To discover the molecular identities of SBARRO cells, we first distinguished single-cell RNA libraries from background by leveraging properties of both single-cell RNA and VBC data from individual experiments. Specifically, using total single-cell RNA data, we identified cell profiles 1) exclusively associated with cell barcodes provided by 10x genomics (corresponding to v2 or v3 chemistry) and exhibiting 2) large UMI counts and low fractions of mitochondrial and ribosomal transcripts, as described previously^30^. In parallel, we used the mutation-collapsed VBC data (see above) to filter and retain those cell profiles with at least a single VBC ascertained with >= 3 (v2 chemistry) or >= 5 (v3 chemistry) UMI counts. We used the union of cell barcodes identified by RNA-based and VBC-based methods to generate digital gene expression matrices (DGEs) for each experiment^56^. DGEs were input into a two-staged analysis pipeline based on independent components analysis (ICA)^30^, a semi-supervised approach for grouping scRNA profiles into clusters then subclusters. scRNA profiles corresponding to cell-cell doublets and cell outliers were identified, flagged and excluded from downstream analyses as described previously^30^. The identities of clusters and subclusters were systematically annotated based on molecular marker expression^30, 57^. Prior to ICA analysis, DGEs were pruned of 1) genes present on the rabies virus or mitochondrial genomes and 2) small scRNA profiles (profiles with >= 500 UMIs (SCC Experiment) or >= 50 genes (noRG Experiment) were retained) to promote high-quality clustering based on host cell nuclear gene expression. Additional DGEs (subject to the same gene and cell filtering criteria) were also generated while including rabies virus genes to aid in the downstream analyses of rabies virus expression:

| Experiment | 10x | FACS | Reads (x1000) |  |  | UMIs (x1000) |  |  | Genes (x1000) |  |  |
| --- | --- | --- | --- | --- | --- | --- | --- | --- | --- | --- | --- |
|  |  |  | Range | Median | Mean | Range | Median | Mean | Range | Median | Mean |
| No RG | V2 | No | 3.9-805.5 | 41.7 | 81.5 | 1.5-92.3 | 7.3 | 12.7 | 0.17-9.5 | 2.9 | 3.5 |
| SCC (SBARRO) | V3 | Yes | 0.6-5957.8 | 30.2 | 53 | 501-291 | 11.1 | 17.6 | 0.049-11.7 | 3.8 | 3.8 |
| SCC (Control) | V3 | No | 1.2-833.2 | 41.2 | 54.7 | 0.97-159.9 | 13.3 | 18.1 | 0.29-11.6 | 4.5 | 4.6 |

To enhance molecular identification of cells in the SCC experiments, an integrated analysis of SBARRO and control cell libraries was performed using LIGER^58^. Control cells were sampled from total cell suspension not subject to FACS and derived from physically adjacent culture wells seeded with the same cell suspensions as SBARRO experiments. Input DGEs for LIGER analysis lacked rabies virus and mitochondrial genes and were filtered to remove “cell-cell doublet” or “outlier” RNA profiles as identified by upstream ICA-based analysis. LIGER alignment and clustering (factorization, k=40, lambda=3; quantile alignment, resolution= 0.4, knn_k=20) results were visualized with UMAP embedding and systematically annotated using marker gene expression. Of the 130.5K SBARRO scRNA profiles, 28.4K were grouped into three clusters (cluster 1, 8 and 13) which contained cells of multiple classes and were defined by expression signatures related to GO biological processes such as “cytokine- mediated signaling pathway (GO:0019221)” (cluster 13; adjusted p value < 2.1×10^-14^) or “PERK-mediated unfolded protein response (GO:0036499)” (cluster 8; adjusted p value < 0.001). To clarify the molecular identities of these 28.4K cells, we re-aligned these SBARRO libraries to the control cells from SCC7 and SCC8 using LIGER (factorization, k=60, lambda=3; quantile alignment, resolution= 0.4, knn_k=20). The resulting analysis split SBARRO libraries across 32 (of 37 total) clusters which exhibited molecular marker expression consistent with known cell populations. Clusters were then systematically annotated in a manner consistent with the initial LIGER analysis guided by the original and re-aligned cluster identities of control cells. In total, we identified n= 20 “granular” cell populations which could be grouped into n=11 “coarse” populations. To identify and visualize genes differentially expressed across granular populations, we used the FindMarkers() and DotPlot() functions from Seurat ^59, 60^. To evaluate which cell populations were sensitive or recalcitrant to rabies virus infection, for each population, we performed a chi-square test comparing the number of SBARRO and control scRNA profiles to the dataset totals and used the resulting residuals as a metric of enrichment or depletion. To determine the relationship between the SBARRO molecular identities and cortical neuron types from the adult mouse cortex, we used LIGER to jointly analyze scRNA profiles from 78.8K uninfected SBARRO control cells and 57K cells from various neocortical regions ascertained by the Allen Institute^18^. We conducted separate analyses for glutamatergic and GABAergic neurons using the same LIGER parameters (factorization, k=10, lambda=30; quantile alignment, resolution= 0.2, knn_k=400).

### Transcriptional identification of starter and presynaptic cells

Starter cells are functionalized after Cre-mediated recombination inverts TVA-mCherry and rabies virus B19G transgenes within the FLEX rAAV genome into the sense orientation with respect to the CAG promoter^61, 62^. RNA-based identification of starter cells in a direct and qualitative manner is complicated using these vectors in the context of 3’ scRNA-seq since 1) low expression caused Cre mRNAs not to be captured with high-probability in Cre+ cells (caused by the limiting MOI of the Syn1-EBFP-Cre rAAV (∼ 8×10^-3^- 8×10^-5^); mild RNA Polymerase II recruitment with the *Synapsin1* promoter; and lack of 3’ motifs to promote mRNA stability) and 2) the vast majority of scRNA-seq reads that align to FLEX rAAV genome do so in the 3’ UTR, a region unaffected by recombination. To overcome these limitations and identify starter cells from scRNA profiles alone, we developed a protocol to selectively amplify and sequence only TVA- mCherry mRNAs transcribed from Cre-recombined rAAV genomes (see *“Selective mRNA Adaptor PCR”* above) and a downstream informatic approach using these data to identify rare, candidate starter cells. Specifically, we developed a binomial test with experiment-specific success rate parameter (updated using an expectation-maximization-like approach) to identify cells in which both recombined TVA-mCherry UMIs (relative to total rAAV UMIs) and total rAAV UMIs (relative to host cell RNA UMIs) were enriched in a manner unlikely to be due to chance (Bonferroni-corrected p < 0.01). Properties of RNA expression that distinguish identified starter cells from presynaptic cells in SBARRO experiments – such as the ratio of recombined TVA- mCherry UMIs / total rAAV UMIs – were observed in scRNA-seq libraries in which starter cells were physically separated from presynaptic cells using FACS.

### Cell-type-specific synaptic network inference using rabies virus barcodes

To facilitate interactive analysis and discovery of cell-type-specific SBARRO networks, we created an R Shiny program (“Terminal E”) which allows dynamic filtering and plotting of VBCs and VBC-based synaptic networks. Terminal E integrates data from VBC libraries and individual SBARRO experiments (organized into “collections” that enable cross-experiment meta-analyses). Library-level data include VBC abundances and user-defined, library-specific lists of single VBCs or VBC pairs to exclude from network inference. SBARRO-level data center around the properties of each cell in each experiment, which, at minimum, include 1) VBC UMI counts; 2) molecular cell type identities; and 3. starter or presynaptic assignments. Inferred synaptic networks are collections of cells which share one or more VBC which are statistically likely to have entered those cells through clonal replication and spread from a single starter cell infection. Terminal E supports the inference of such networks through two stages of VBC filtering. During the first stage, filters completely exclude VBCs from network consideration. We only considered VBCs with >= 7 UMIs (SCC experiments, v3 10x chemistry) or >= 3 UMIs (No RG Experiments, v2 10x chemistry). We additionally removed those VBCs with high abundances in genomes of the infecting library (FI trust score >= 5; see below). We further identified specific VBCs for exclusion by 1) comparing their presence and abundance in rabies virus libraries to behavior transducing 17K starter cell scRNA profiles (containing 28.8K founder infections) or 2) by comparing across n=23 independent SBARRO experiments. We excluded the named categories of VBCs below based on the following criteria (Extended Data Fig. 6b):

1. “Felony” VBCs (n=234). Absent from library but observed in > 1 of 5,015 starter cells (Fig. 2f).
2. “Misdemeanor” VBCs (n=78). Present in library, but observed in > 2 starter scRNA profiles with library frequency < 10^-6^ or observed in > 8 starter scRNA profiles with library frequency < 10^-5.5^ (Fig. 2f).
3. “Cross Experiment” VBCs (n=239) Absent from library but observed in > 1 of 23 independent SBARRO experiments.

In the second stage, VBCs or VBC sets are filtered such that those retained were suitably rare enough to enter the experiment through a single starter cell. Critical to this stage of experiment-specific VBC filtering is an estimate of the total number of experiment-specific founder infections starter cells; a subset of which lead to cell-to- cell spread (spreading founder infections). To estimate the number of spreading founder infections for each experiment, we developed an analytical approached designed to mimic founder infections by drawing samples of VBCs from the viral library. We created distributions describing the number of library draws (n=10 replicates, with VBC replacement) required to match the number of unique VBCs observed in >=2 cells present in each experiment; median values set the experiment- specific founder infection estimates. To evaluate whether each VBC or VBC set was suitably rare enough to be included for network inference, we calculated and assigned an “founder infection trust” (FI trust score). The “FI trust” score equates to the number of spreading founder infections that could in theory occur for that VBC or VBC set before a second founder infection was expected (at a given probability) by leveraging library VBC frequencies:

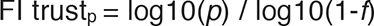

Where *f* is the frequency of an individual VBC in the library (or, for the VBC set, the inferred frequency calculated by multiplying the frequencies of each VBC member) and *p* is the probability of avoiding a second occurrence of the VBC or VBC set. For a given experiment, VBC or VBC sets that were retained to infer synaptic networks were those for which FI trustp > experiment estimates for founder infections. For SCC experiments, *p* = 0.9. Inferred networks containing a single scRNA profile assigned as a starter cell were then split into postsynaptic (i.e. the starter) and presynaptic cells; networks lacking a starter cell were starter-orphaned and all scRNA profiles were presumed presynaptic. Terminal E facilitates the comparisons of presynaptic network size (the number of presynaptic cells) and cell type composition across postsynaptic starter cells of different types. After first stage filtering, VBCs absent from the library were presumed to be rare and assigned the lowest library frequency value to match library VBCs counted with a single UMI. After stratifying presynaptic networks by starter cell type, presynaptic cell type compositions were compared using a Chi- Square Test. Presynaptic network sizes were assigned to each scRNA profile using described rules (Extended Data Fig. 6c) and presynaptic network sizes were compared across starter cell types using a Wilcoxon Rank-Sum test.

### Postsynaptic RNAs associated with the number of presynaptic partner cells

Postsynaptic RNA profiles from the SCC experiments identified by as one of four abundant starter cell types (glutamatergic neurons, interneurons, SPNs or astrocytes; n=144) were binned based on their inferred presynaptic network size: small (2-4 cells); medium (5-6 cells) or large (7+ cells). These bins approximate inflections in the total distribution of presynaptic network sizes (Figure 4b). To determine if viral load was associated with presynaptic network size category, the fraction of total cellular mRNA derived from all five rabies virus genes was compared using a Wilcoxon Test. A similar strategy was used to test for differences in innate immune responses, using an aggregate expression score derived from the 564 genes (a subset of 646 total genes for which we detected a transcript) curated as part of the mouse innate immune response^63^. To identify candidate gene expression differences associated with presynaptic network size, we compared starter cell RNA profiles associated with “small” and “large” presynaptic across cell subtypes rather than types, to avoid confounds due to differences in subtype compositions. Because these comparisons involved small numbers scRNA profiles (range: 2 to 28 scRNA profiles; mean, 6.3; median, 4) we used the following strategy to identify genes and contextualize how likely such differences were likely to arise by chance. First, we used Fisher’s Exact Test to evaluate differences in UMI counts across scRNA profiles aggregated by small or large presynaptic size category. We considered only those genes with >= 25 UMIs thus lessening the burden of multiple hypothesis testing and corrected our p value cut < 0.05 by the number of tests completed within each subtype comparison. Second, for the genes which passed threshold, we used a Wilcoxon Test to determine whether normalized RNA levels (removing contributions from rabies virus mRNAs before normalization) differed across the population of individual cells associated with each presynaptic network size category, using p < 0.05 as our cutoff. To help determine which of the genes we identified were likely due to chance, we repeated this testing procedure for 100 permuted comparisons in which each starter cell RNA profile was replaced by a presynaptic cell RNA profile of the same subtype choose at random. Genes identified by multiple permuted replicates in the same cell subtype were flagged as potentially spurious and not considered further. Log fold expression changes were calculated after normalizing the number of gene-specific UMIs by total UMIs for large and small presynaptic network categories and then scaling the data to 100,000 transcripts after the addition of a pseudocount.

### Identifying mRNAs correlated with rabies virus transmission across SPN development

Monocle3^24, 64^ was used to calculate pseudotime scores for scRNA profiles of developing SPNs, from neural precursor cells to mature neurons, after preprocessing (method=PCA; number of dimensions = 10) and alignment SBARRO and control libraries (alignment_k=3000). To identify modules of genes with similar expression levels over pseudotime, all genes were first tested for pseudotime- associated expression using the graph_test() function Genes with q values = 0 were retained and modules identified using the find_gene_modules() function with resolution = 0.001. Pseudotime-ordered cells were grouped into 10 bins. For each bin, 1) the fraction of total cells that were SBARRO (rather than control) in origin were calculated and 2) a meta-control cell was created by summing control cell UMIs and normalizing such that mRNA expression values summed to 100,000. Pearson correlations were calculated for each detected gene by comparing SBARRO fractions and normalized gene expression values across the pseudotime bins. SBARRO- correlated genes (n=3,309) were defined as those genes in which r >= 0.75. SBARRO-correlated genes were tested for gene set enrichment using PantherGO^25, 26, 65^ via EnrichR^66^ and SynGO^29^ and compared to two sets (n= 1000 replicate gene selections/set) of n=3,309 control genes, selected either 1) from expression-matched deciles built from the developmentally mature SPN meta-control cell (bin = 10) or from 2) at random from genes with expressed RNA.

## EXTENDED DATA FIGURES

**Extended Data Figure 1.**
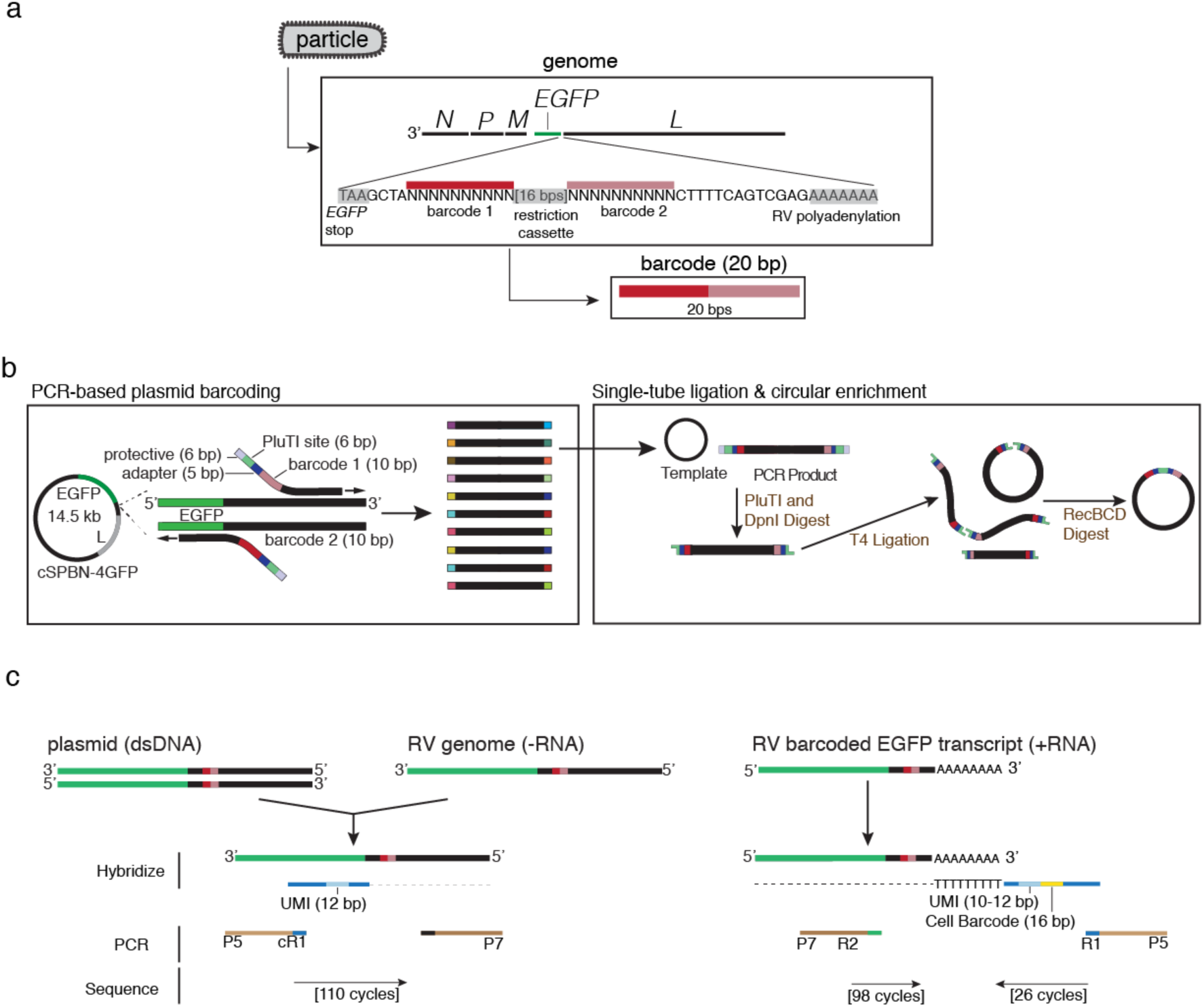
Molecular workflows for PCR-based plasmid barcoding and UMI-based VBC quantification across plasmids, anti-sense rabies virus genomes and mRNAs. **a.** Schematic showing the VBC cassette integrated into the rabies genome. The 20 bp VBC consists of two 10 bp barcodes linked by a restriction cassette and sits 13 bps away from the rabies polyadenylation signal. **b.** Diagram describing the novel PCR-based strategy for barcoding DNA plasmids as applied to the cDNA version of the SAD-B19 RV*dG*-EGFP genome (pSPBN-4GFP). Forward and reverse primers containing 10 bp randomers - tailed by 17 bps of containing adaptor sequence, the PluTI site and protective bases – are targeted to adjacent regions of the template plasmid. Amplification results in a collection of linear dsDNA molecules each of which contain a unique combination of terminal 10 bp barcodes, which are then circularized and enriched (versus both template plasmid and remaining linear products) through a single-pot reaction. **c.** Illumina sequencing-based strategy for quantifying plasmid and anti-sense genomic barcodes (*left*) or barcoded *EGFP* mRNA (*right*) using unique molecular identifiers (UMIs). UMI-containing oligonucleotides (UMI = 12 bp randomer) with a shared PCR handle are hybridized adjacent to and then polymerized through the barcode cassette on ssDNA or anti-sense RNA genomes. The UMI-tagged ssDNA molecules are then selectively PCR amplified using primers that contain Illumina P5 and P7 sites and sequenced on an Illumina flow cell such that 110 Read 1 cycles cover the barcode cassette. Barcoded *EGFP* mRNA is selectively amplified from sc- cDNA using P5 and P7 containing primers and sequenced on an Illumina flowcell such that 26-28 Read 1 cycles that capture the 16 bp Cell Barcode and 10-12 bp UMI introduced by 10x Chromium v2 or v3 chemistry and 98 Read 2 cycles which extend through the barcode cassette (**Methods)**.

**Extended Data Figure 2.**
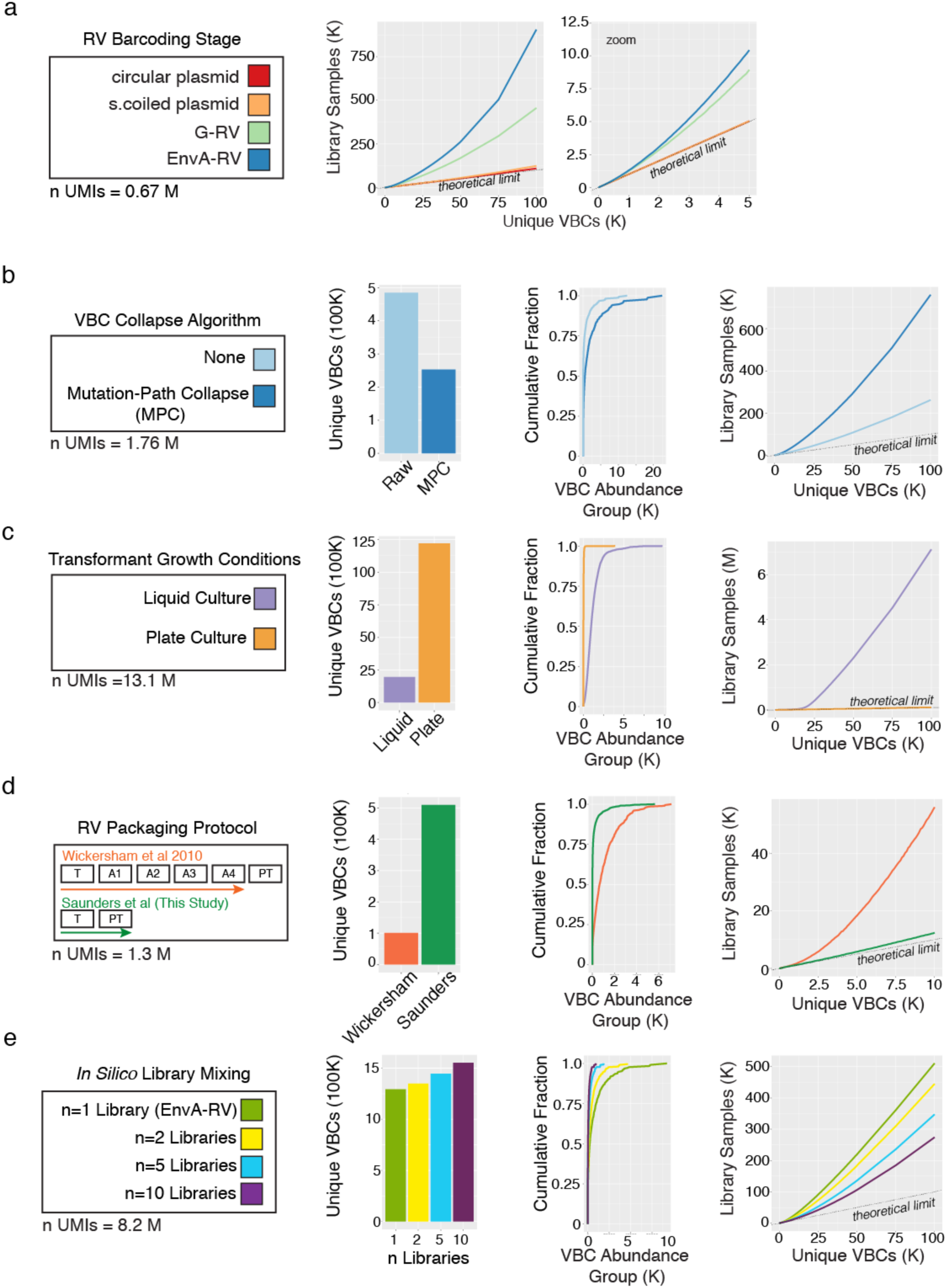
Accurate and systematic quantification of VBC abundances guides optimized protocols for plasmid barcoding and barcoded rabies virus packaging. **a.** Longitudinal assessment of VBC diversity across each stage of rabies virus packaging protocol, as assayed through the library sampling procedure in which total library samples are plotted against the number of unique VBCs ascertained from each sample. (Companion data to Fig. 1d**,e**. Plot includes data from Fig. 1f along with additional conditions). The dotted line shows maximum theoretical diversity (in which every drawn VBC is unique). **b-e**. Quantification of VBC abundance and diversity across various protocol conditions (sampled with equivalent UMIs, *far left*) by plotting (*from left to right*) total unique VBCs; cumulative distribution of UMIs by abundance group (as in Fig. 1e); and number of unique VBCs ascertained from a given number of library samples (as in **a** above). **b.** The effect of “mutation- path collapse” (MPC), an informatic approach implemented to help account for artifactual inflation of barcodes driven by mutations to barcode sequences incurred during library amplification or sequencing (**Methods**). **c.** The effect of *E.coli* growth conditions (plated or liquid culture) after transformation with circular barcoded plasmid library. Barcodes were sampled from super-coiled plasmid DNA. **d.** The effect of rabies virus packaging protocols, comparing the widely used Wickersham et al. 2010 protocol versus the barcode diversity optimized protocol reported in this study (Saunders et al. 2021). Barcodes were sampled from anti-sense genomes extracted from EnvA-pseudotyped libraries. **e**. The effect of combining different numbers of independent and equivalently diverse barcoded EnvA-pseudotyped libraries *in silico* (**Methods**).

**Extended Data Figure 3.**
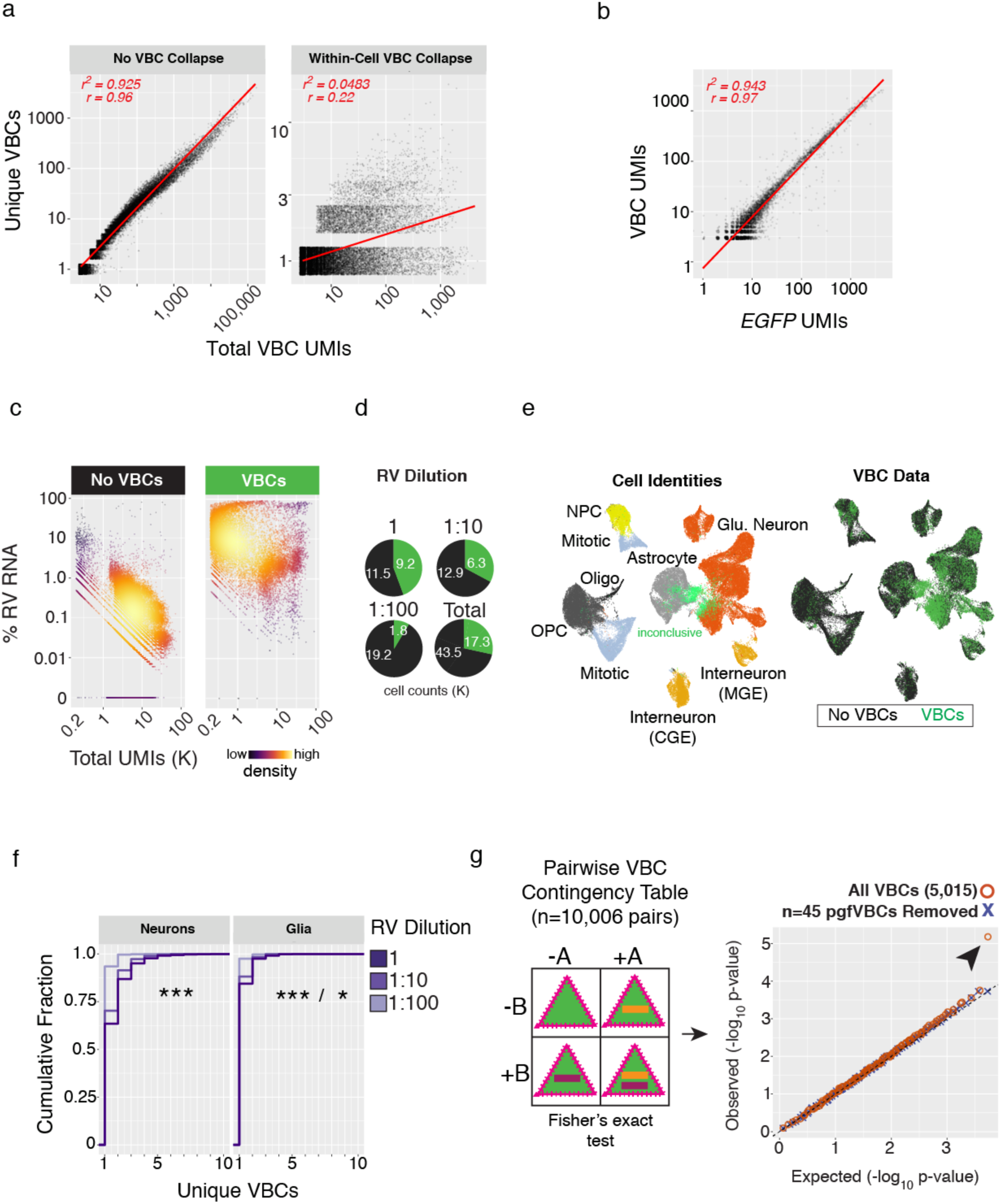
Integrating host cell RNA and viral VBC data for thousands of individual starter cells relate how properties of barcoded rabies libraries behave across founder infections resolved by cell type. **a.** An informatic approach (“Within-Cell VBC Collapse”, **Methods**) for reconstructing accurate VBC sequences and UMI counts from scRNA data in light of amplification and sequencing artifacts. *Left*, without correction, mutations in barcode sequences incurred during PCR and Illumina sequencing inflate the number of unique VBC sequences observed in each scRNA profile in proportion to the number of total VBC UMIs, leading to a strong correlation (r = 0.96). *Right*, following “Within-Cell VBC Collapse,” the relationship between unique VBCs and total VBC UMIs becomes more independent (r = 0.22). **b.** Single cell UMI counts for VBCs (inferred from 3’ *EGFP* UTR sequencing) and the *EGFP* mRNAs (inferred from host cell RNA sequencing) are highly correlated (r = 0.97), indicating a strong correspondence across independent sequencing datasets (data are from a single SBARRO experiment, n = 6,979 cells). **c-e.** Single-cell RNA profiles ascertained from brain cells grown *in vitro* expressing *TVA* but not rabies *G* were transduced with EnvA-RV*dG*-*EGFP*VBC at three different concentrations (no dilution, ”1” (MOI ∼ 15); diluted one in ten “1:10” (MOI ∼ 1.5); or one in a hundred “1:100” (MOI ∼ 0.15); **Methods**). **c.** VBC data are comprehensively ascertained from infected (“VBCs”; n = 17,283) but not uninfected (“No VBCs”; n= 43,533) scRNA profiles over a wide range of UMI counts and percentages of total viral RNA. **d.** Pie charts illustrating percentages of scRNA profiles for which VBC data were ascertained (green) or not ascertained (black) across rabies virus dilution conditions. Total cell counts are listed. **e.** UMAP embedding of 60,816 scRNA profiles color-coded by molecular identity (*left*) or VBC ascertainment status (*right*) following LIGER analysis (**Methods**). A subset of infected scRNA profiles (n = 2,635) could not be definitively identified (light green). **f.** Cumulative distribution of unique VBCs per cell, grouped by neuron versus glia type and color-coded by rabies virus dilution. Increasing rabies virus titer leads to more unique VBCs per cell, but does so in a sublinear manner with respect to MOI, suggesting an intrinsic, cell-type-specific limit to the number of independent founder infections (* = p < 0.05; *** = p < 0.001, Kolmogorov–Smirnov Test). **g.** Testing VBC independence in the context of 17.2K starter cell founder infections with multiple VBCs. *Left*, a schematic of the contingency table comparing the number of scRNA profiles in which two VBCs (“A”, purple; “B”, orange) occur together (+A/+B), independently (+A or +B) , or are not observed (-A/-B). VBC pairs which occurred together more than chance (n=45 of 10,009 total pairs with Bonferroni-corrected p < 0.05, Fisher’s Exact test) were considered putative genome fusion VBCs and flagged. *Right*, Q-Q plot comparing observed vs expected p-values - expected p-values were generated after VBCs were randomized across scRNA profiles - when all VBCs pairs were considered (orange circles) and after n=45 pgfVBC pairs were removed (purple “x”). Arrowhead highlights the inflation of observed p-values away from the expectation of random driven by putative genome fusion VBCs (**Methods**).

**Extended Data Figure 4.**
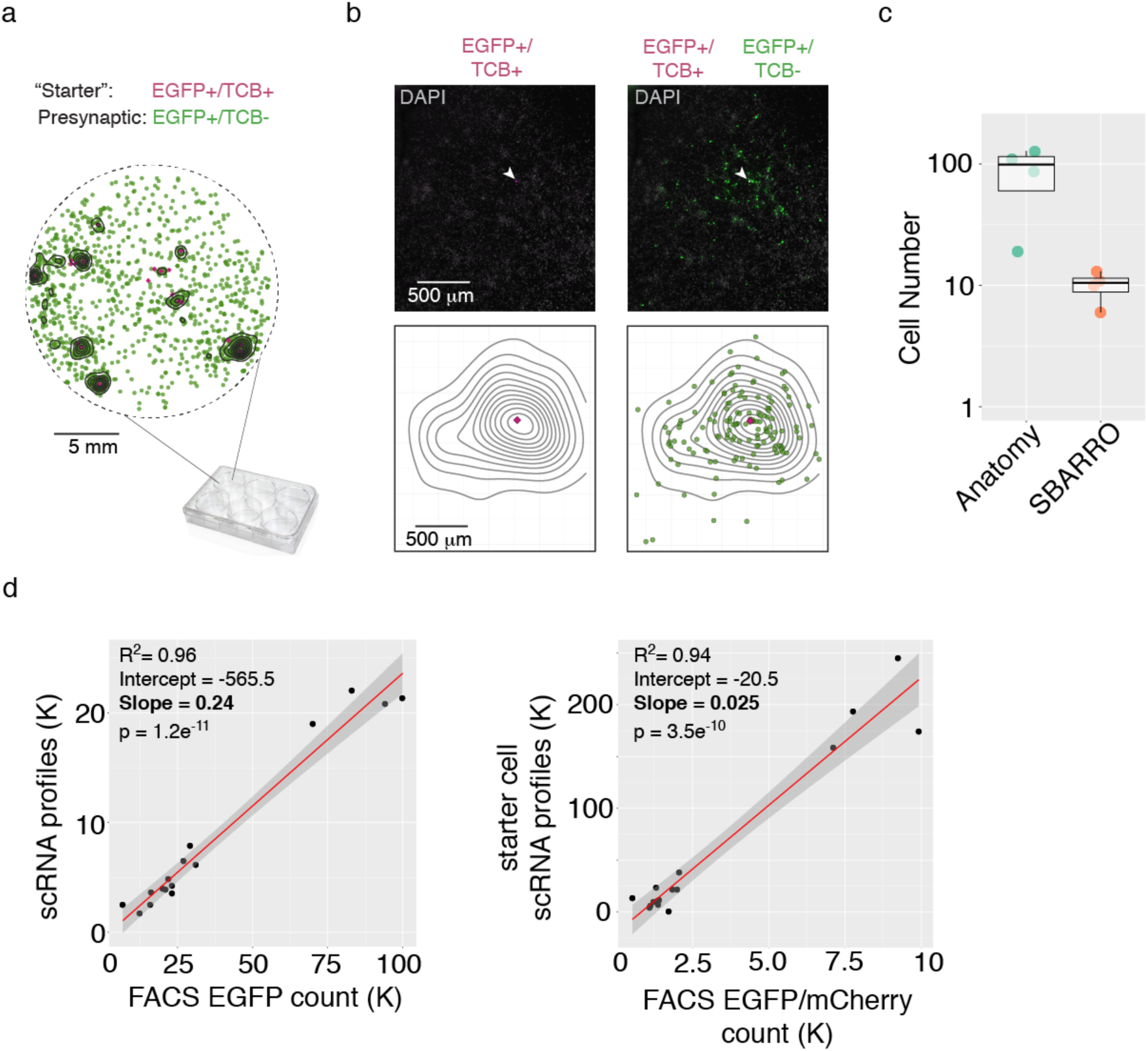
Anatomy of monosynaptic rabies virus spread *in vitro*. **a-b.** Monosynaptic cell-to-cell spread events of rabies virus in cell cultures derived from dissociated embryonic mouse cortex exhibit stereotyped spatial patterning. In each culture well, a small subset of potential starter cells was endowed using a rAAV Cre-recombinase based strategy followed by transduction of EnvA-RV*dG*-*EGFP*VBC. Fluorescent scans of whole culture wells distinguish the locations of these spatially sparse starter (EGFP+/TVA-mCherry+, magenta) and presynaptic cells (EGFP+/TVA- mCherry-, green) (**Methods**). **a.** Locations of starter and presynaptic cells derived from a scan of a representative culture well. Presynaptic cells tend to spatially cluster around starter cells, but are also observed at in a distributed fashion at greater distances from any individual starter. Contours illustrate areas density of rabies virus infected cells. **b.** Higher magnification view showing the locations of a single starter cell (*left*) and presynaptic cells in close proximity (*right*). *Top*, fluorescent images. *Bottom*, plot of extracted cell locations. **c.** A comparison of inferred presynaptic network sizes based on anatomical imaging (the number of clustered presynaptic cells) or SBARRO sequencing (the number of presynaptic scRNA profiles based on uCIPs). Experiments were performed in parallel from neighboring culture wells grown from the same cell suspension. The largest four inferred networks from each modality are shown. **d.** A comparison of FACS-based cell counts and scRNA profiles for all SBARRO cells (*left*) or just starter cells (*right*) fit with linear models for which the slope was used to estimate the sample rate (n=16 culture wells; **Methods**). **d.** A comparison of FACS-based single-cell RNA sampling rates of rabies virus infected EGFP+ cells (n=16 culture wells). A linear model of slope = 0.24 described the relationship (R^2^ = 0.96 and p = -1.2e-11), suggesting scRNA was sampled from 24% of FACS-enriched cells (**Methods**).

**Extended Data Figure 5.**
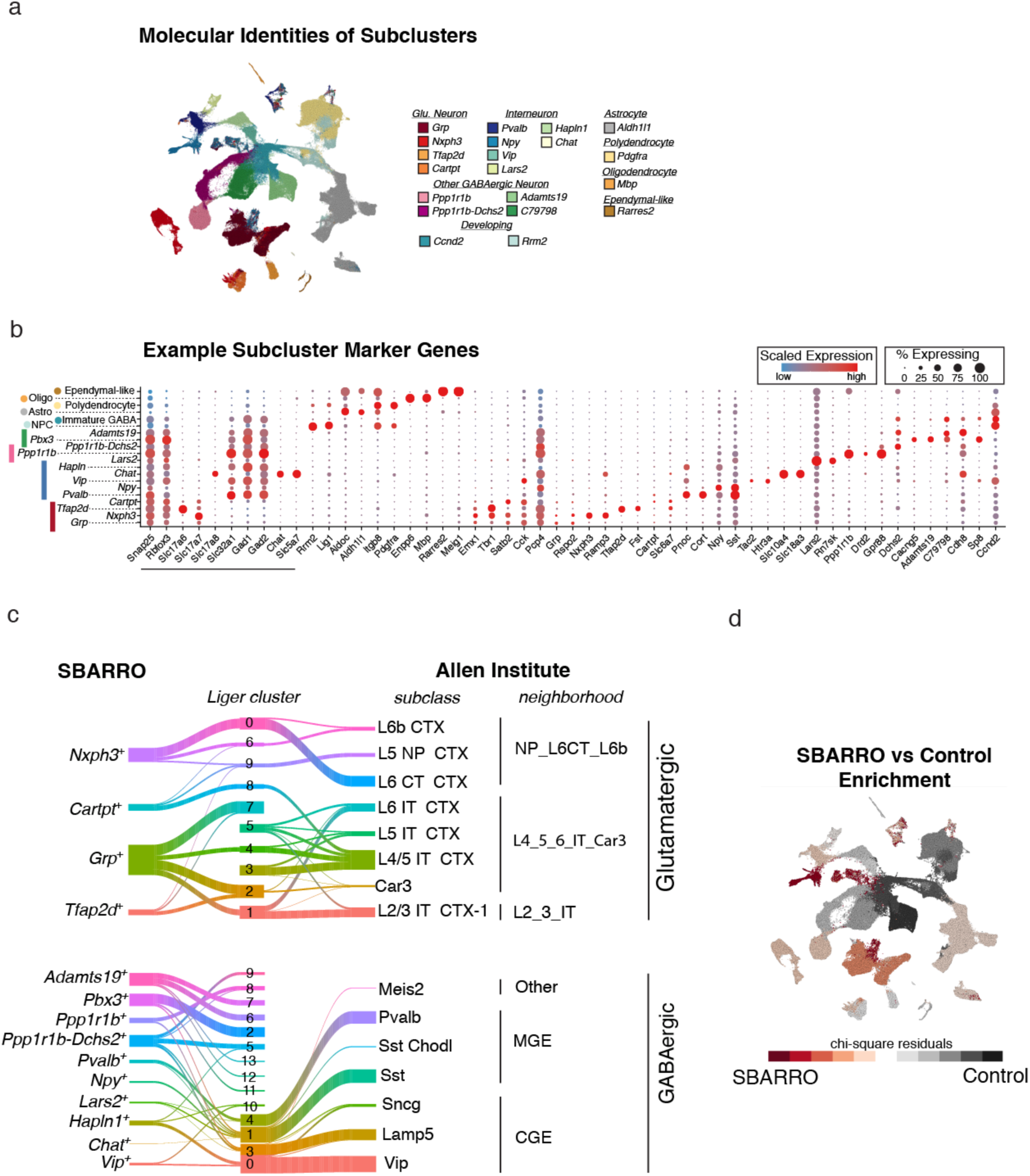
Assigning molecular identities to SBARRO scRNA profiles. **a.** UMAP embedding color-coded by granular molecular subtypes (subclusters). **b.** Dotplot of example marker gene expression patterns across subcluster populations. Common markers for neurons and neuron types are underlined; additional genes pairs were selected for each population based on differential expression analysis (**Methods**). **c.** Sankey plots showing molecular homologies between scRNA profiles from SBARRO control cells *in vitro* and adult mouse cortex *in vivo*^18^ following LIGER analysis of glutamatergic (SBARRO, n = 24,155 profiles; Allen Institute, n= 38,899) and GABAergic (SBARRO, n = 54,713 profiles; Allen Institute, n= 18,163) neurons (**Methods**). **d.** Quantifying enrichment or depletion of SBARRO libraries as compared to control scRNA profiles. Color-code shows chi-square residuals after the number of SBARRO/control RNA profiles within each coarse molecular population are compared to dataset totals (**Methods**).

**Extended Data Figure 6.**
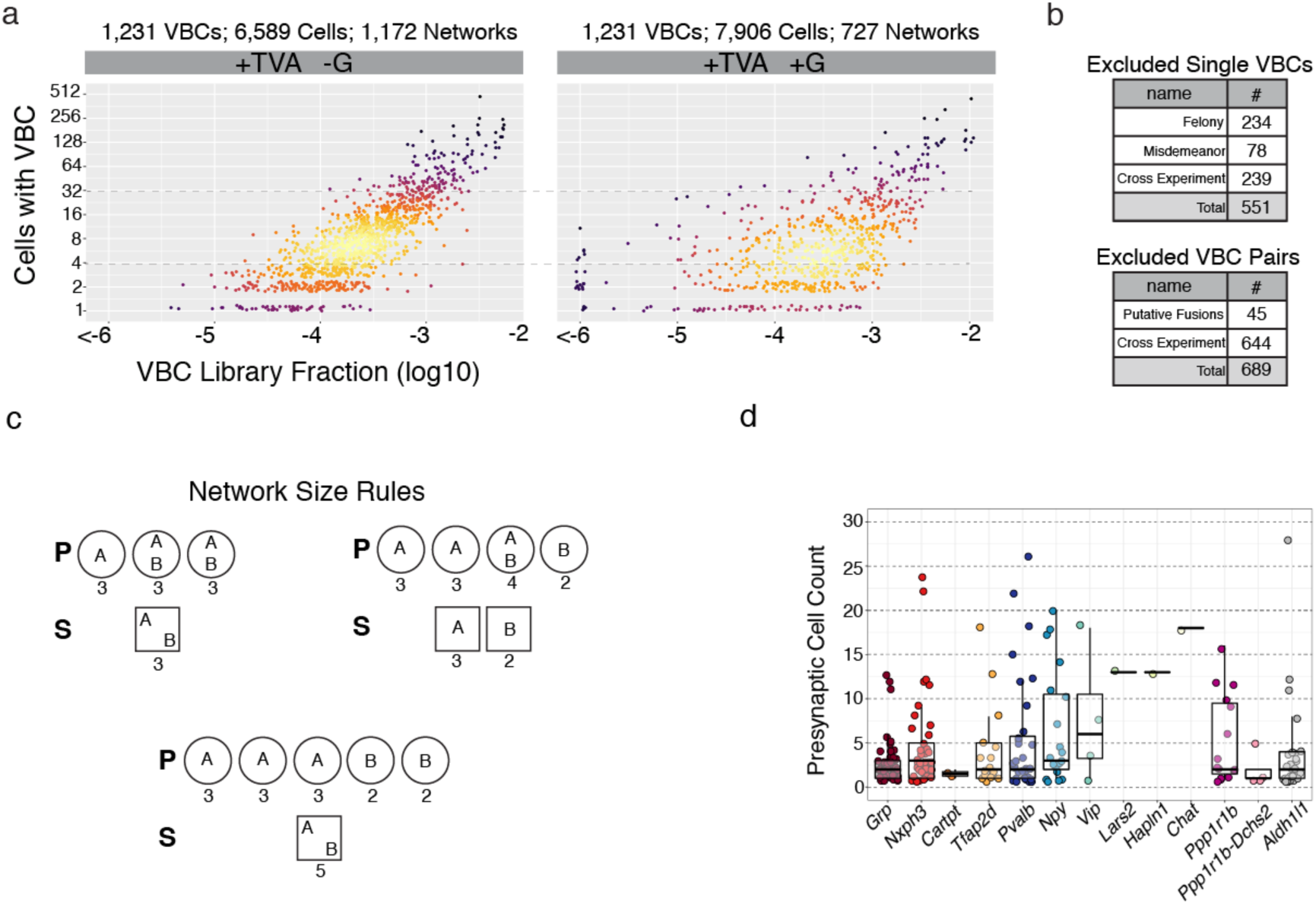
SBARRO inference of synaptic networks through VBC- based CIPs. **a.** The effect of rabies virus spread on the relationship between VBC library abundance and number of cells in which each VBC was ascertained. *Right*, a single experiment (“SCC07_1e3_A”) in which EnvA-RV*dG*-*EGFP*VBC founder infections were complemented with glycoprotein endowing monosynaptic retrograde spread (+G). *Left*, a version of the G- starter cell corpus (Fig. 2f), consisting exclusively of founder infections, randomly down-sampled to match equivalent VBC numbers (n=1,231). **b.** Tables of VBCs (*top*) and VBC pairs (*bottom*) excluded from network inference using EnvA-RV*dG*-*EGFP*VBC (**Methods**). **c.** Schematic describing how the “Network Size” parameter was calculated for starter cells (squares) and presynaptic cells (circles) based on examples using two VBCS, “A” and “B”. Network Size values are listed below each cell. **d.** Inferred presynaptic network sizes by starter cell subtype for all identified starter cells.

**Extended Data Figure 7.**
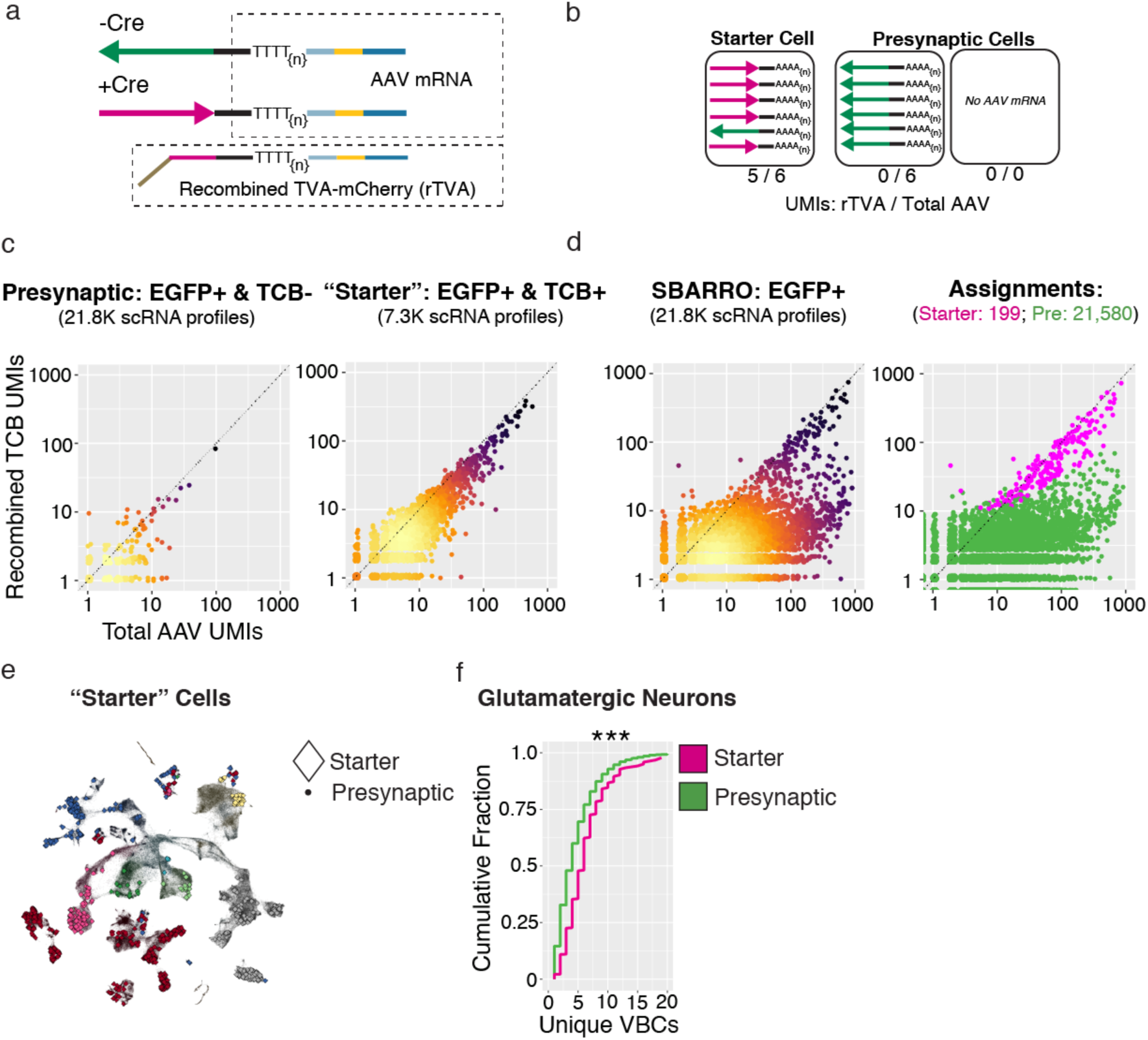
Assigning starter cell identities to SBARRO scRNA profiles. **a-c.** Identifying starter cells from SBARRO scRNA profiles. Starter cells are endowed with functional *TVA-mCherry* and *G* mRNAs after Cre-mediated recombination of CAG-Flex-TVA-mCherry and CAG-Flex-B19(G) rAAV genomes. **a.** Schematic describing the 3’ end structure of recombined (+Cre, magenta) or unrecombined (-Cre, green) rAAV mRNAs (such as those encoding *TVA-mCherry* and *G*) after single-cell barcoding and first strand synthesis. The sequenced region critical to determining the identity of the expressed gene – the black bar in the dashed upper box – is unaffected by recombination, thus the vast majority of single cell counts of rAAV mRNAs enabled by standard 3’ scRNA-seq (i.e. Drop-seq, inDrop and 10x) are not recombination-informative. To generate sequencing libraries selectively for recombined rAAV mRNAs, we amplified and independently sequenced (using Illumina flowcells) only recombined *TVA-mCherry* transcripts (bottom dashed box; **Methods**). **b**. Cartoon illustrating recombined vs unrecombined rAAV mRNA content of starter and presynaptic cells. Starter cell RNA profiles are enriched for recombined rAAV mRNAs, while presynaptic cells are enriched for unrecombined molecules or have no detectable rAAV mRNA counts. **c**. Validation of rAAV mRNA signatures for starter and presynaptic cells after physical separation via FACS. Scatter plots comparing UMI counts of recombined *TVA-mCherry* mRNA vs Total rAAV mRNA for scRNA profiles resulting from FACS-based separation and independent library generation of presynaptic (*left*, EGFP+/TVA-mCherry-) or starter (*right*, EGFP+/TVA-mCherry+) cells. Points falling along the dotted unity line are cell profiles for which all rAAV mRNAs are from recombined *TVA-mCherry* transcripts. scRNA profiles from sorted starter cells have higher counts of Total rAAV mRNAs and those counts are largely from recombined *TVA-mCherry mRNAs*. Lighter colors indicate higher point densities. **d.** Assigning starter cell identities to SBARRO scRNA profiles (n=21,580) from a single experiment (“SCC07_1e2_C”). *Left*, scatterplot of recombined *TVA-mCherry* vs Total rAAV mRNA counts. *Right*, color-coded by starter (n=199) or presynaptic (n=21,580) assignment based on the results of a binomial testing in which starter RNA profiles exhibit statistical enrichments for recombined *TVA-mCherry* counts (versus Total rAAV counts) and Total rAAV counts (versus all UMIs; **Methods**). **e.** UMAP locations of starter and presynaptic cells. **f.** Cumulative distribution of unique VBCs across glutamatergic neuron starter and presynaptic cells (*** = p < 2.2e-16, Kolmogorov–Smirnov Test).

**Extended Data Figure 8.**
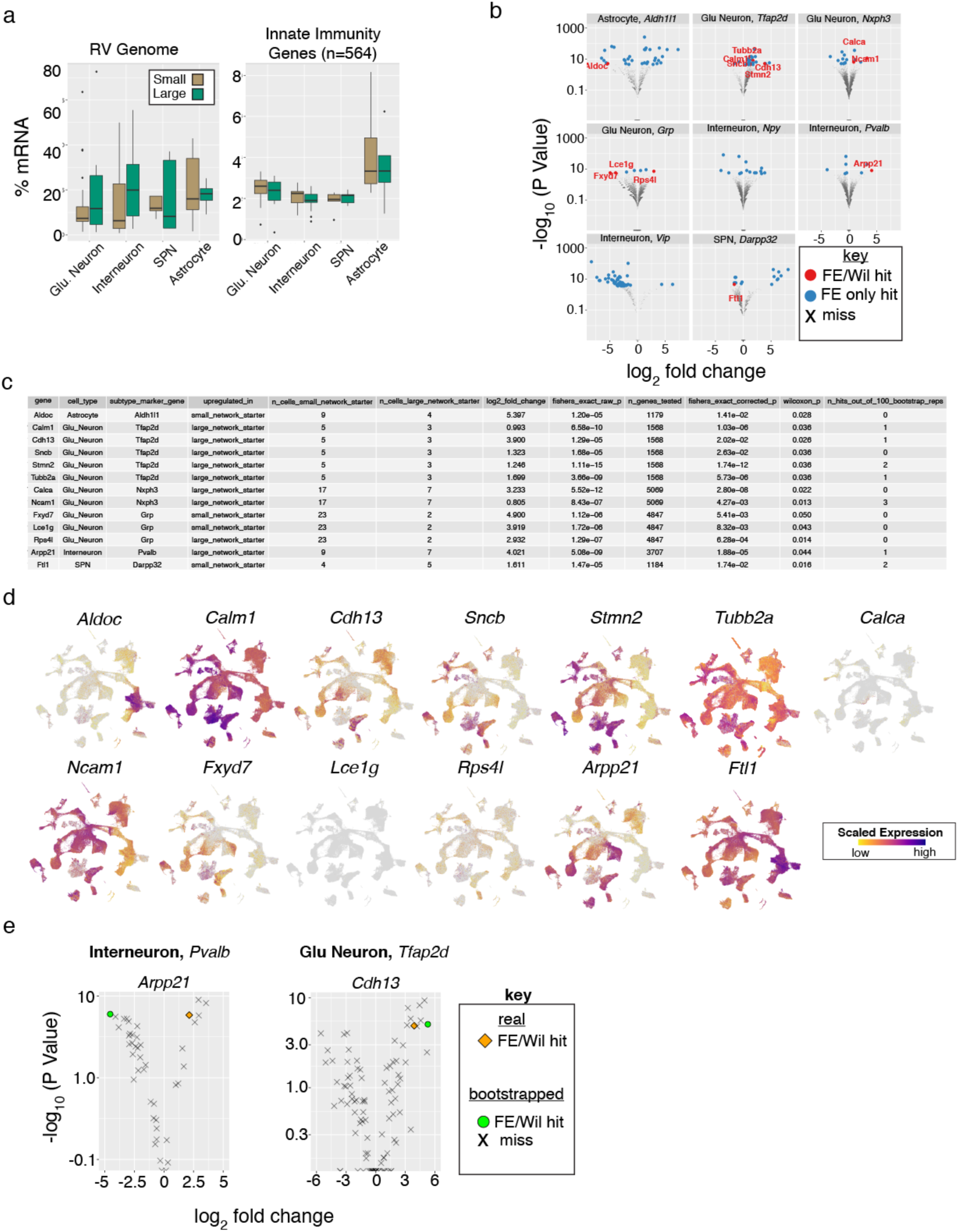
Properties of postsynaptic starter cell infection and host cell RNAs associated with presynaptic network size inferences. **a.** Viral load (% of all mRNAs derived from 5 rabies virus genes; *left*) and innate immunity expression scores (aggregated from 584 curated genes^22^; *right*) across starter cell RNA profiles (n=144) did not show detectable differences across “large” or “small” presynaptic network size groupings for four major brain cell types (p > 0.05, Wilcoxon Test). **b.** Volcano plots illustrating results from differential expression testing of starter cell RNA profiles comparing “large” and “small” presynaptic network size categories by starter cell subtypes. UMI counts for each gene of sufficient expression were aggregated by inferred presynaptic network size category then compared (Fisher’s Exact Test; **Methods**). Genes passing corrected p value thresholds (p < 0.05, blue dots) were further tested for differences in single-cell scaled expression (Wilcoxon Test; **Methods**). Those genes that pass this additional test (p < 0.05) were considered hits and labeled (red dots). **c**. Summary table describing differential expression results for those genes identified in b. **d**. Expression plots for the genes identified in c. **e**. Volcano plots comparing differential expression results for *Arpp21* in *Pvalb*+ Interneurons (*left*) and *Cdh13* in *Tfap2d*+ Glutamatergic Neurons (*right*) in real data or 100 permuted replicates in which starter cell RNA profiles were randomly replaced by presynaptic profiles of the same subtype (**Methods**). The real data is shown with a gold diamond; permuted replicates passing aggregate UMI (Fisher’s Exact Test) and scaled expression comparisons (Wilcoxon Test) are shown as green circles; all other comparisons are shown with grey crosses.

**Extended Data Figure 9.**
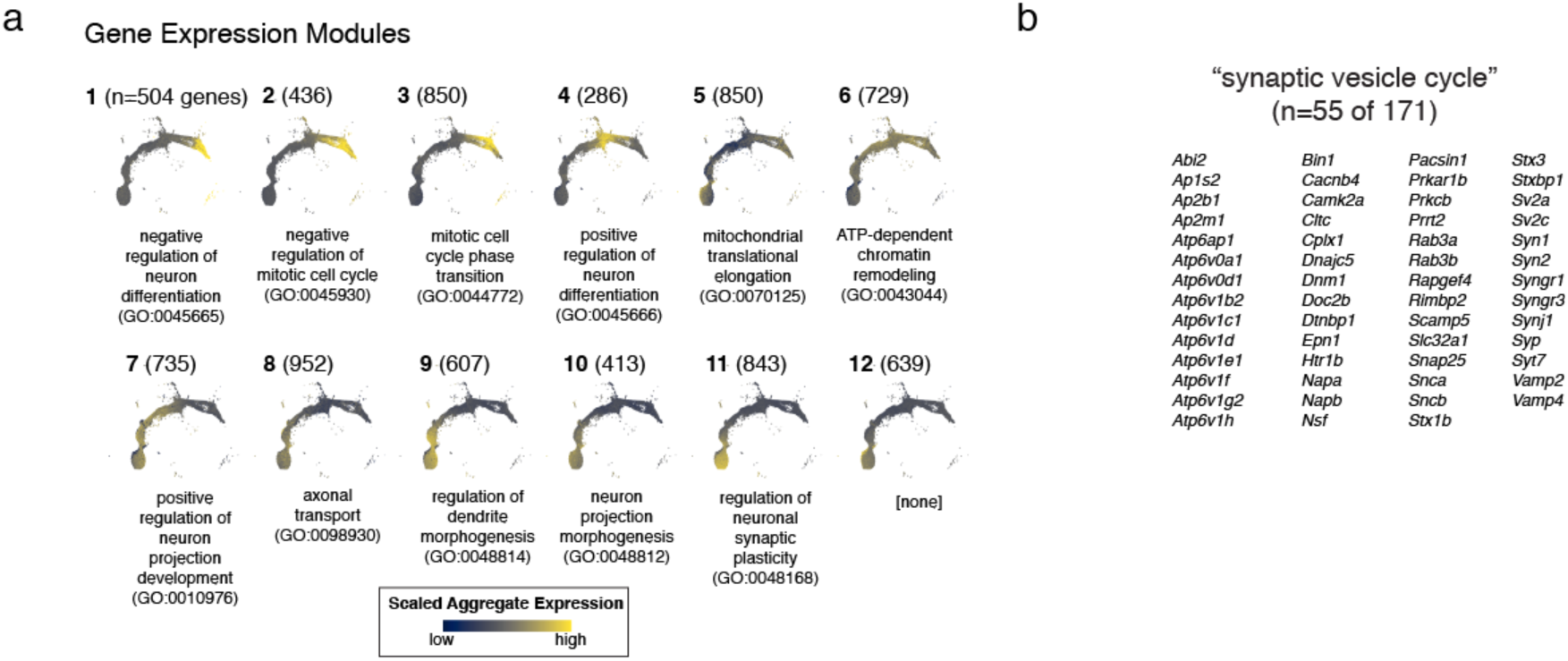
Molecular correlates of rabies virus transmission during SPN development. **a.** Modules of gene expression (n=12) identified with Monocle3 (**Methods**). For each numbered module, the count of associated genes is shown parenthetically, module expression is color-coded by aggregate expression and a representative enrichment for biological process gene ontologies categories is shown (adjusted p < 0.05). **b.** Names of rabies virus-transmission correlated “synaptic vesicle cycle” genes (n=55 of 171 in SynGO category GO:0099504).

---

**Supplementary Table 1.**
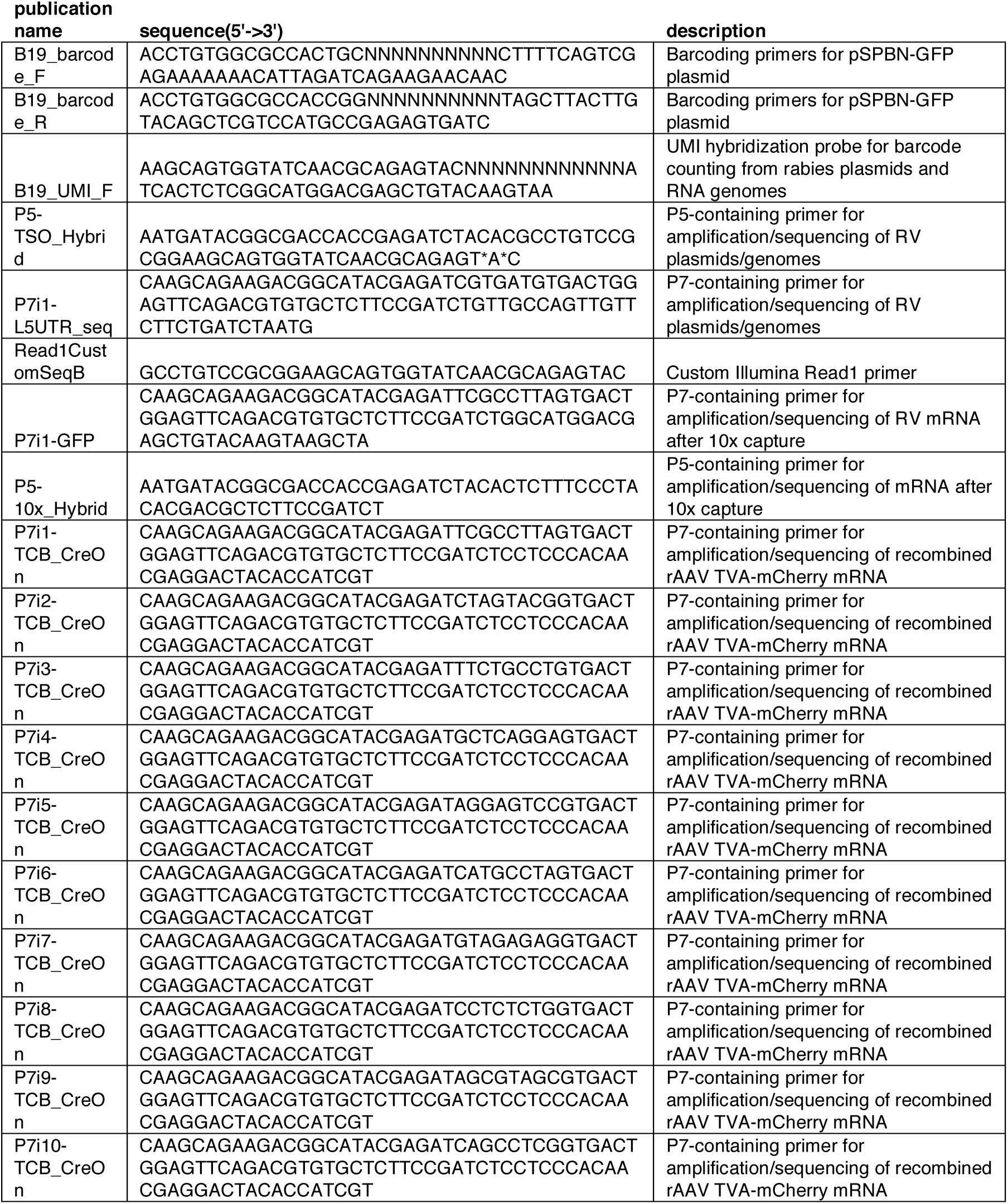
Oligonucleotide Guide

